# From homeostasis to credit assignment: a signed-XOR connectomic motif for local directional error signalling

**DOI:** 10.64898/2026.06.05.730322

**Authors:** María Peña Fernández, Alejandro González Ríos, Lara Lloret Iglesias, Jesús Marco de Lucas

## Abstract

Biological neural circuits are widely thought to require local error signals that tell synapses not only that a prediction is wrong, but also in which direction to change. We previously proposed that a six-neuron XOR motif acts as a homeostatic comparator: matched sensory and predictive signals cancel locally, whereas mismatches propagate an error signal. We also showed that a shallow autoencoder can learn MNIST using a signed-XOR learning rule with local decoder errors and random feedback alignment, without gradient backpropagation. Here we introduce the *signed-XOR motif*, an eight-neuron, twelve-edge directed signed circuit that extends the XOR comparator with two feedback channels of opposite neurotransmitter identity. By construction, the motif can convert a binary mismatch into directional error signalling, with one pathway encoding potentiation and the other depression, while respecting Dale’s principle. We provide open-source tools to enumerate the motif at connectome scale and test its enrichment against degree- and sign-preserving null models. The motif is enriched 24.3× in *C. elegans* (*Z* = 52.2), significantly enriched in 59/80 FlyWire *Drosophila* neuropils including AVLP_L (13.9×, *Z* = 94.4), and strongly enriched in layers 2/3–5 of a biophysically detailed mouse primary visual cortex model (global 315×; per-pivot medians up to 852 ×) while absent from layer 6. The same layer-specific pattern is found in the axon-proofread subset of the EM-reconstructed MICrONS connectome. A Brian2 leaky integrate-and-fire implementation reproduces the signed-XOR truth table, remains robust to Poisson drive, produces a graded signed error, and requires a fast-spiking parvalbumin-like pivot. These results identify signed-XOR as a recurrent connectomic pattern compatible with local homeostatic error cancellation and directional credit-assignment signals.

**Author Summary:** How does a brain decide which of its connections to adjust when it makes a mistake? Unlike an artificial network, it has no global error signal supplied from outside: each connection can react only to the neurons it directly touches. We ask whether a small, repeating wiring pattern could provide such a local correction signal. The pattern we study, the signed-XOR motif, compares an incoming signal with the brain’s own prediction of it. When the two agree, the circuit stays quiet, so already-expected activity is not relayed onward. When they disagree, it does more than flag an error: it also indicates the direction of the fix, routing it through two separate channels, one meaning “strengthen”, the other “weaken”, consistent with the biological rule that each neuron acts with a single sign. We provide open software to search for this pattern in three nervous systems, a worm, a fly, and a detailed model of mouse visual cortex, and find it more often than chance wiring predicts, with a striking layer-specific distribution in cortex. We also simulated the eight-cell circuit with realistic spiking neurons and confirmed that it can perform the computation, but only when its inhibitory cell is a fast-spiking type like those concentrated in the enriched layers. We do not claim that any brain uses this circuit to learn or memorize. What we provide is a specific motif that could deliver a local, directional error signal that may be useful for a neuromorphic implementation.

## 1 Introduction

### 1.1 Credit assignment in biological networks

How does a brain decide which synapse to strengthen? In artificial neural networks, the answer is given by a global loss function ℒ and the gradient *∂*ℒ*/∂w* delivered by backpropagation [1]. In biological networks the same question has no consensual answer: backpropagation in its classical form is widely considered biologically implausible [2, 3], yet brains evidently learn. A number of plausible alternatives have been proposed: random feedback alignment [4], predictive-coding hierarchies [5], burst-mediated plasticity [6], target propagation, and equilibrium propagation, among others. What unites these proposals is their reliance on a *local error signal* carried by the network itself rather than computed from a centralised loss function [7].

A local error signal has two requirements. First, it must be *local*: the synapse should be able to read it from neurons it directly contacts. Second, it must be *informative*: it should tell the synapse not only that there is an error, but how to correct it, in particular in which direction (potentiation versus depression) to update the weight. The first requirement constrains the topology of the circuit that delivers the signal; the second constrains its semantics.

### 1.2 The XOR motif as a binary mismatch detector

In a previous note [8], we proposed that a six-neuron motif organised around a single inhibitory hub could implement an exclusive-OR (XOR) comparison between two binary signals, e.g. an external sensory input *x* and an internal prediction *f* (*x*). The motif’s output neuron fires whenever *x* ≠ *f* (*x*) and remains silent when they match, providing a discrete, locally-readable mismatch detector. We also described how such motif could be embedded in a computational neural network based in the Liquid Time Constant model, providing a basic learning for binary sequences. In another note [9] we showed how the motif appeared to be quite popular in three different connectomes that were openly available: *C. elegans, Drosophila* and mice.

### 1.3 The missing piece: directional feedback

In a following contribution [10], we extended the application of the XOR motif embedding it as a per-output-bit comparator in a one-hidden-layer autoencoder trained on MNIST, and we found that it converged to 97.7% bit-wise reconstruction accuracy in five epochs without any backpropagation or global error computation, and that an unsupervised latent representation supported 89.8% digit classification by a subsequent linear readout.

The bare XOR motif was, however, only half of what was needed for this result. The Hebbian-style update rule used was:

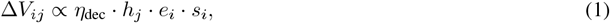

where *h*_*j*_ is the hidden-unit activation, *e*_*i*_ is the XOR error bit, and *s*_*i*_ = 2*x*_*i*_ − 1 ∈ { − 1, +1} is the *sign* of the desired correction (potentiate when the output undershoots the target, depress when it overshoots), depends on a signed quantity. The bare XOR motif emits only *e*_*i*_ ∈ {0, 1}. The sign *s*_*i*_ must come from somewhere else.

Within a circuit obeying Dale’s principle (the empirical regularity that a single neuron releases a single neurotransmitter species and therefore exerts a single sign of postsynaptic effect [11]), the natural way to deliver a signed signal locally is to split it across two pathways: an excitatory aggregator that fires when potentiation is required, and an inhibitory aggregator that fires when depression is required. We call this enriched topology the *signed-XOR motif*. It augments the six-node XOR core with two additional neurons, F^+^ (excitatory identity) and F^−^ (inhibitory identity), and four feedback edges (Section 2, Figure 1).

**Figure 1:**
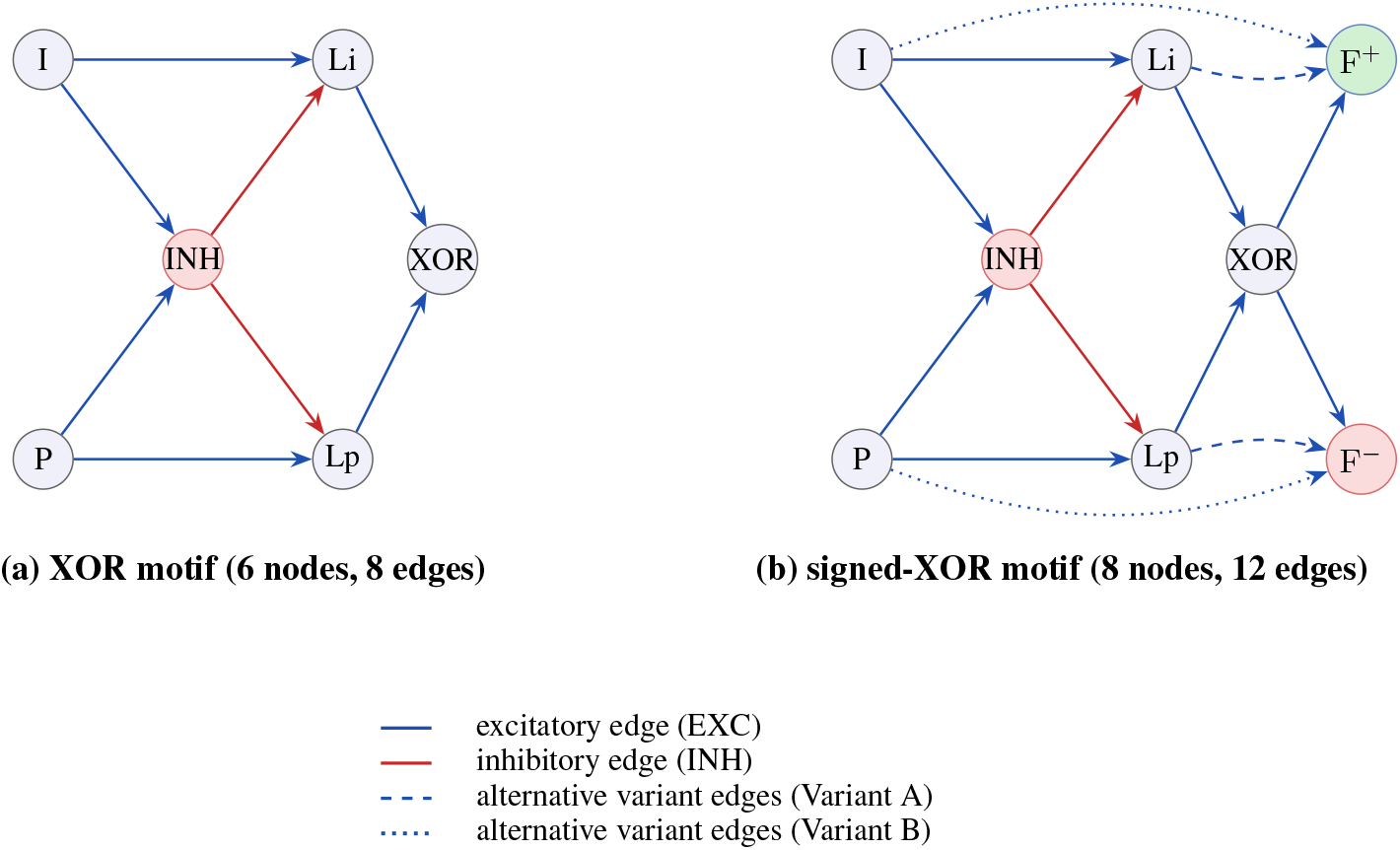
The XOR and signed-XOR motifs. *(a)* The bare XOR motif: six neurons, eight directed signed edges, with an inhibitory hub (node INH) gating the convergent excitation onto the XOR node, which is active iff exactly one of the two inputs (nodes I, O) is active. *(b)* The signed-XOR motif extends (a) with two feedback aggregators of opposite neurotransmitter identity (F^+^, F^−^) and four edges. The shared edges *XOR* → F^+^ and *XOR* → F^−^ inherit XOR node’s sign. Variant A (dotted: *I* → F^+^, *P* → F^−^) provides input-side feedback; Variant B (dashed: *Li* → F^+^, *Lp* → F^−^) provides intermediate-side feedback.

### 1.4 Hypothesis and contribution

If signed-XOR-like circuits are useful substrates for local directional error signalling, then real connectomes should bear a statistical signature: the motif should be present at a higher frequency than expected under a null model that preserves basic network statistics (degree distributions and neurotransmitter composition) but disrupts higher-order organisation. Conversely, if the signed-XOR is merely an arbitrary eight-node subgraph among many, it should appear at chance levels.

This study tests that hypothesis across three connectomic systems. Our contributions are:

1. We **formalise** the signed-XOR motif as an 8-node, 12-edge directed signed graph with explicit identity constraints on each node (Section 2).
2. We provide **open-source C tools** that enumerate the motif and a reference Python pipeline, validated to within a single mapping against an original NetworkX implementation (Sections 3 and D; [12]).
3. We **enumerate** the motif in the *C. elegans* connectome [13] (Section A), in the FlyWire *Drosophila* connectome [11, 14] across 80 neuropils (Section B), and in a biophysically detailed mouse V1 model [15] (Section C), also corroborated in EM-reconstructed MICrONS connectome [16].
4. We **compare** the three species (Section 4.5), showing that the signed-XOR motif is over-represented relative to degree- and sign-preserving nulls in the invertebrate connectomes and the V1 model, and is anatomically structured — layer-specific — in mouse cortex.
5. We discuss the mechanistic implication: real connectomes contain more instances of this directional-feedback template than expected under sign- and degree-preserving null models, as would be expected if such circuits were useful substrates for local learning (Section 6).

## 2 The signed-XOR motif: definition and mechanistic role

### 2.1 Formal definition

We define the signed-XOR motif as a directed signed subgraph on 8 distinct nodes labelled {*I, P, Li, Lp, INH, XOR*, F^+^, F^−^} with 12 directed edges (Figure 1b). Each edge carries a sign in EXC, INH inherited from the neurotransmitter identity of its source neuron. The required edges are:

- **XOR core** (8 edges):

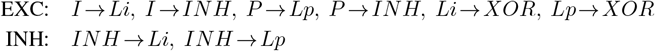
- **Feedback shared edges** (2 edges, both with sign equal to the neurotransmitter identity of *XOR* node):

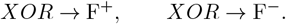
- **Variant A** OR **Variant B** (2 edges):

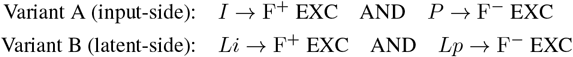

A motif instance is counted whenever Variant A holds, Variant B holds, or both hold simultaneously (Variant AB).

#### Identity constraints

The two feedback aggregators have asymmetric neurotransmitter requirements: F^+^ must be excitatory and F^−^ inhibitory. Node XOR may be either, but the two shared feedback edges from it must match its sign. Roles *I, Li, P, Lp* (the XOR-core afferents) must be excitatory; role INH (the inhibitory hub) must be inhibitory.

#### Distinctness

All 8 neurons must be pairwise distinct.

#### Symmetry

The bare XOR core admits the 2-fold automorphism *I* ↔ *P, Li* ↔ *Lp*. In the signed-XOR motif this would-be symmetry would also require F^+^ ↔ F^−^, which violates the asymmetric identity constraint and is therefore not an automorphism. As a consequence, each biological signed-XOR instance is found exactly once; the raw mapping count and the distinct biological count coincide. (For the bare XOR motif, in contrast, raw = 2× distinct.)

### 2.2 Mechanistic interpretation

The role of node XOR in the bare motif is the binary mismatch indicator *e* ∈ {0, 1}. The role of F^+^ and F^−^ is to deliver the *sign* of the correction. In the previously cited autoencoder framework [10], the synaptic update of Equation (1) requires both the error magnitude *e* (node 5) and its direction *s* (under-vs. over-shoot). Within a Dale-respecting circuit, a natural way to encode a signed quantity is by splitting it across two pathways of opposite neurotransmitter identity: F^+^ active means potentiate; F^−^ active means depress. Both aggregators receive identical input from node XOR (the shared edges), ensuring they fire only when the mismatch is present; the variant-specific edges gate *which* sign of correction is appropriate. Variant A reads the sign from the input neurons; Variant B from the intermediate XOR-internal neurons. Both implement the same logical function but make different commitments about where the sign-defining information is read out.

### 2.3 Homeostatic interpretation and excitation–inhibition balance

The XOR core can be read as a local homeostatic comparator. When the two compared signals match, either no excitatory lane is driven, or both lanes are driven and the inhibitory pivot vetoes their relays; in both cases the comparator remains silent. Matched activity is therefore cancelled locally rather than amplified downstream. When the two signals differ, exactly one relay escapes the veto and the circuit emits a mismatch signal. The signed extension preserves this stopping rule while splitting the mismatch into two correction channels, *F* ^+^ and *F* ^−^.

This homeostatic interpretation is related to excitation–inhibition balance, but it should not be read as a derivation of a precise anatomical ratio. In the excitatory-XOR instantiation used in the spiking validation, six of the eight roles are excitatory and two inhibitory; in the more general connectomic enumeration, the XOR readout itself may have either sign. The relevant point is therefore qualitative: a small inhibitory component gates several excitatory pathways to prevent the propagation of already-predicted activity, while allowing unpredicted activity to generate a directional error signal. This is reminiscent of the broad cortical design principle that excitation is numerically dominant but locally controlled by inhibitory interneurons, often approximated by a 4:1 excitatory-to-inhibitory ratio at the population level.

### 2.4 Rationale and testable hypothesis

A signed-XOR-like circuit is not the only way a brain could implement directional feedback. Burst-dependent plasticity [6] delivers a graded multiplicative signal via a single feedback pathway; predictive-coding architectures [5] embed signed errors in paired prediction and reconstruction neurons. The hypothesis we test is more modest: *if* biological circuits implement localised signed-error signals at the level of single-neuron-resolution motifs, then a topology that respects Dale’s principle and provides a sign-decoded readout is a plausible candidate. Under this hypothesis, signed-XOR-like motifs should occur more frequently than expected by chance in appropriately controlled connectomic null models.

## 3 Methods

We summarise the shared methodology here; per-species data sources, parameters and full result tables are given in the Appendix, in Section A (*C. elegans*), Section B (*Drosophila*) and Section C (mouse V1), and the null model is detailed in Section D. All code is open-source [12].

### 3.1 Input format and edge sign

Each connectome is reduced to a directed, signed edge list: per connection, a pre-synaptic and a post-synaptic neuron and an excitatory/inhibitory label. The sign is derived per dataset (from a curated per-edge classification in *C. elegans*, from predicted neurotransmitter in *Drosophila* and from cell-type identity in mouse V1), with the species-specific rules and their biological justification given in the respective appendix.

### 3.2 Motif counting

A general-purpose subgraph-isomorphism search (e.g. NetworkX) is prohibitive at connectome scale. We use a hand-tuned C counter that pivots on the most-constraining node (the inhibitory hub, role 6, with ≥ 2 outgoing inhibitory edges), then iterates over ordered pairs of its inhibitory targets, locates the excitatory predecessors, and identifies the convergent target; the signed-XOR counter adds two loops over candidate (F^+^, F^−^) aggregators. Two modes are supported. **Embedded** (non-induced): an instance is counted if the required edges are present, regardless of additional edges among the motif’s nodes, the appropriate measure of a motif embedded in a larger circuit. **Induced**: the induced subgraph on the motif’s nodes must match the motif’s edge set exactly. The counter was validated against an independent NetworkX pipeline on AVLP_L to the unit (Section B).

### 3.3 Null model

We compare observed counts against a degree-preserving null ensemble generated by the unbiased edge-swap MCMC (Markov Chain Monte Carlo) of Roberts and Coolen [17], applied independently within each sign (or neurotransmitter) layer so that every neuron’s in- and out-degree of each sign is preserved exactly. This preserves Dale’s principle while randomising higher-order structure. Default mixing is *T* = 100 proposed swaps per edge per layer. Implementation details, the per-species layering choice, and a data-integrity safeguard are provided in Section D.

### 3.4 Significance

For each (network, motif, mode) we generate *N* replicates, count the motif in each, and report the real count *C*_obs_, the null mean 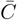 and SD *σ*_*C*_, the *Z*-score 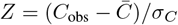, the fold enrichment 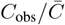, and the empirical two-sided *p*-value

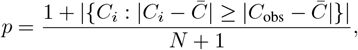

with floor 1*/*(*N* + 1). For the *Drosophila* multi-neuropil scan we additionally report Benjamini–Hochberg FDR-corrected *q*-values.

## 4 Results

### 4.1 Overview across three species

We enumerated the bare XOR and the signed-XOR motif in three connectomes: the *C. elegans* hermaphrodite connectome (whole organism), the FlyWire *Drosophila* brain (80 neuropils analysed separately), and a biophysically detailed model of mouse V1 (locally, around inhibitory pivots). The signed-XOR motif, counted in embedded mode, is significantly over-represented in all three (Table 1). Per-species detail, parameters and full tables are in Sections A to C; below we state the headline results and then synthesise.

**Table 1.** Embedded signed-XOR motif over-representation across species, *n* = 1000 degree- and sign-preserving null replicates (*N* = 100 per pivot over 60 pivots for the Allen mouse V1 model). The *Drosophila* row is the AVLP_L neuropil; *C. elegans* is whole-organism; the Allen V1 row is the layer-4 `i4Pvalb` cell type (summed real / mean null over its pivots; fold and *Z* are per-pivot medians). The MICrONS row is the layer-4 `i4Pvalb` count on the axon-proofread subset of the EM-reconstructed connectome (1,708 neurons, 10 pivots), with layer assignment taken from the MICrONS cell-type labels. *Folds and Z-scores are not comparable across rows: the substrates differ in reconstruction method, edge-weight semantics, pivot sample size, and null stringency. For MICrONS no fold or Z is reported (entries n/a): on this dense, spatially organised graph a degree- and cell-type-preserving null produces more motifs than the real graph and so does not provide a chance baseline; the MICrONS row is a real-count description, not an enrichment claim; see Section C*. Counts are biological-distinct; all *p* at the resolution floor.

| Organism / region | Scale | Real | Null mean | Fold | $Z$ |
| --- | --- | --- | --- | --- | --- |
| *Drosophila* AVLP_L | whole neuropil | 1,813,119,238 | 130,426,267 | 13.9× | 94.4 |
| *C. elegans* herm. | whole organism | 110,399 | 4,537 | 24.3× | 52.2 |
| Allen V1 model (L4 i4Pvalb) | per pivot type | 11,122,740 | 20,100 | 515.9× | 2663 |
| MICrONS V1 (L4 i4Pvalb) | EM, proofread | 55,870 | n/a | n/a | n/a |

**Table 2.**
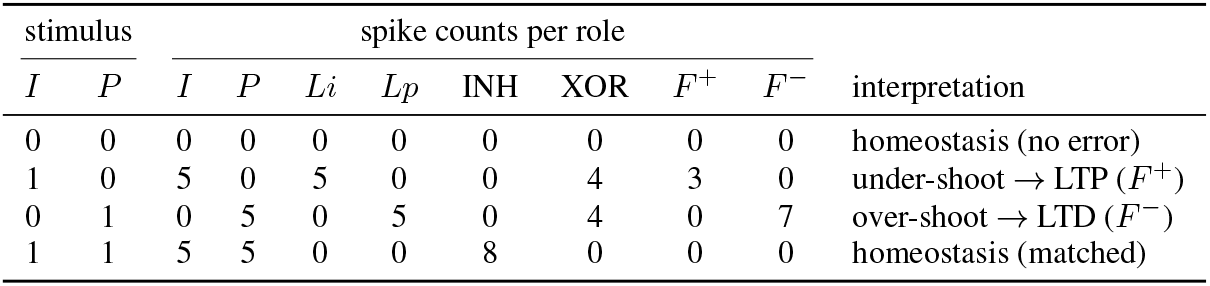
Spike counts per role over a 100 ms trial for each input condition (first two columns are the binary drive to *I* and *P*). A role reads out as active at ≥ 2 spikes. The comparator (XOR) is active only if *I* and *P* differ, and *F* ^+^*/F*^−^ partition the two mismatch directions. The table is reproduced deterministically; under 20 Hz Poisson background (25 seeds) the full table is recovered on 88% of trials.

**Table 3:**
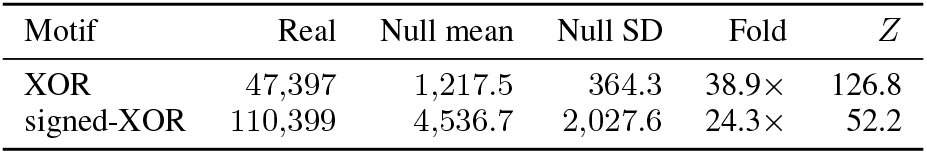
XOR and signed-XOR motifs in the *C. elegans* hermaphrodite connectome, embedded counting, *n* = 1000 degree- and sign-preserving null replicates (sign layered by the curated EXC/INH classification).

### 4.2 *C. elegans*: XOR motifs strongly enriched

In the *C. elegans* hermaphrodite connectome (300 neurons; 2,933 excitatory and 253 inhibitory chemical synapses after self-loop removal; Section A), the embedded signed-XOR motif occurs 110,399 times against a null of 4,536.7 ± 2,027.6 (24.3×, *Z* = 52.2, *p* < 10^−3^, *n* = 1000). The bare XOR motif is even more enriched (47,397 vs. 1,217.5 ± 364.3; 38.9×, *Z* = 126.8). Both were counted in embedded mode (induced counting is uninformative on this small, densely clustered graph; Section A). In the nematode connectome the XOR-type convergent excitatory/inhibitory connectivity is broadly observed, not only its feedback-completed form, we may conjecture if this is due to the direct connection from its neurons to other cell types, that may play such feedback role.

### 4.3 *Drosophila*: the signed-feedback extension is enriched beyond the XOR core

On the AVLP_L neuropil (the reference region, *n* = 1000 nulls; full 2 × 2 matrix in Table 4), the embedded signed-XOR motif is enriched 13.9× (*Z* = 94.4), whereas the bare XOR core on the same data and in the same mode is enriched only 4.2× (*Z* = 60.9). Because both counts would scale linearly with the underlying connectivity if the feedback edges were incidental, the ~3.3-fold larger enrichment of the signed motif indicates that the feedback-completed extension carries additional structure beyond what the XOR core alone predicts. Extending the embedded signed-XOR scan to all 80 neuropils (Section B, Table 6), the motif is significant after Benjamini–Hochberg FDR correction in 59*/*80 neuropils, with extreme enrichments in GNG (6,675×), SAD (5,481×), WED_R (1,725×) and the inferior-posterior slope IPS_R (1,274×), and significant *depletion* only in the antennal lobes (AL_L, AL_R) and CRE_R. The bare XOR scan, computed in the strict induced mode (Table 5), shows the opposite aggregate sign (depletion, 0.56×): this is the expected dense-network signature of induced counting (Section 6.3) and is *not* comparable mode-for-mode with the embedded signed scan. The rigorous within-species XOR-vs-signed comparison is the embedded 2 × 2 of Table 4.

**Table 4:** XOR and signed-XOR motifs in FlyWire *Drosophila* AVLP_L, *n* = 1000 replicates, mixing *T* = 100, per-neurotransmitter randomisation. Real counts are biological-distinct.

| Motif | Mode | Real | Null mean | Fold | $Z$ |
| --- | --- | --- | --- | --- | --- |
| XOR | embedded | $4.88 \times 10^8$ | $1.15 \times 10^8$ | **$4.23\times$** | **60.9** |
| XOR | induced | $1.05 \times 10^6$ | $3.43 \times 10^6$ | $0.31\times$ | $-11.8$ |
| signed-XOR | embedded | $1.81 \times 10^9$ | $1.30 \times 10^8$ | **$13.9\times$** | **94.4** |
| signed-XOR | induced | $1.09 \times 10^5$ | $3.27 \times 10^5$ | $0.33\times$ | $-5.9$ |

**Table 5:** Bare XOR motif, induced counting: per-neuropil enrichment in FlyWire *Drosophila, n* = 1000 degree- and Dale-preserving null replicates. ⋆ marks significance at FDR *α* = 0.05 (Benjamini–Hochberg *q*). 30/80 neuropils significant.

| Neuropil | Nodes | Edges | Real | Null (mean ± SD) | $Z$ | Fold | $p$ | $q$ |
| --- | --- | --- | --- | --- | --- | --- | --- | --- |
| AL_L | 3,196 | 36,659 | 40,759 | 34,803.7 ± 3,447.9 | 1.73 | 1.17× | 0.07493 | 0.135 |
| AL_R | 3,392 | 36,826 | 36,989 | 28,904.9 ± 3,076.5 | 2.63 | 1.28× | 0.01099 | 0.0293 $\star$ |
| AME_L | 355 | 1,424 | 3 | 10.4 ± 9.4 | -0.79 | 0.29× | 0.4226 | 0.5366 |
| AME_R | 378 | 1,400 | 0 | 12.6 ± 13.5 | -0.93 | 0.00× | 0.1908 | 0.2726 |
| AMMC_L | 1,376 | 13,398 | 2,575 | 1,686.4 ± 280.4 | 3.17 | 1.53× | 0.003996 | 0.01453 $\star$ |
| AMMC_R | 1,666 | 12,217 | 277 | 281.9 ± 71.5 | -0.07 | 0.98× | 0.9481 | 1 |
| AOTU_L | 1,456 | 12,139 | 5,217 | 4,672.8 ± 1,915.7 | 0.28 | 1.12× | 0.7702 | 0.8837 |
| AOTU_R | 1,649 | 14,440 | 4,851 | 4,809.7 ± 1,850.2 | 0.02 | 1.01× | 0.976 | 1 |
| ATL_L | 1,344 | 6,334 | 22 | 42.8 ± 25.6 | -0.81 | 0.51× | 0.3666 | 0.4808 |
| ATL_R | 1,284 | 5,838 | 27 | 33.0 ± 24.5 | -0.25 | 0.82× | 0.7732 | 0.8837 |
| AVLP_L | 4,493 | 98,418 | 1,049,112 | 3,434,574.6 ± 188,503.1 | -12.65 | 0.31× | 0.000999 | 0.004701 $\star$ |
| AVLP_R | 5,703 | 131,152 | 2,341,688 | 5,998,278.9 ± 319,684.6 | -11.44 | 0.39× | 0.000999 | 0.004701 $\star$ |
| BU_L | 259 | 632 | 0 | 0.1 ± 0.3 | -0.17 | 0.00× | 1 | 1 |
| BU_R | 350 | 1,043 | 0 | 0.0 ± 0.4 | -0.04 | 0.00× | 1 | 1 |
| CAN_L | 537 | 2,997 | 13 | 9.1 ± 9.0 | 0.43 | 1.43× | 0.6673 | 0.8213 |
| CAN_R | 931 | 3,860 | 8 | 2.4 ± 5.7 | 0.97 | 3.27× | 0.08392 | 0.1428 |
| CRE_L | 5,196 | 36,450 | 3,957 | 8,566.2 ± 2,324.0 | -1.98 | 0.46× | 0.03996 | 0.07797 |
| CRE_R | 5,067 | 37,359 | 2,942 | 12,372.7 ± 2,556.7 | -3.69 | 0.24× | 0.004995 | 0.01737 $\star$ |
| EB | 938 | 17,022 | 384 | 4,846.8 ± 730.0 | -6.11 | 0.08× | 0.000999 | 0.004701 $\star$ |
| EPA_L | 1,629 | 11,651 | 311 | 793.9 ± 225.2 | -2.14 | 0.39× | 0.03397 | 0.06967 |
| EPA_R | 1,401 | 9,605 | 280 | 445.6 ± 144.7 | -1.14 | 0.63× | 0.2388 | 0.3351 |
| FB | 3,127 | 49,145 | 94,547 | 61,629.4 ± 4,953.1 | 6.65 | 1.53× | 0.000999 | 0.004701 $\star$ |
| FLA_L | 1,890 | 14,969 | 785 | 366.3 ± 102.4 | 4.09 | 2.14× | 0.003996 | 0.01453 $\star$ |
| FLA_R | 2,060 | 17,993 | 1,594 | 842.8 ± 197.9 | 3.80 | 1.89× | 0.003996 | 0.01453 $\star$ |
| GA_L | 417 | 1,843 | 0 | 40.5 ± 25.3 | -1.60 | 0.00× | 0.07892 | 0.1373 |
| GA_R | 312 | 1,318 | 0 | 24.4 ± 18.5 | -1.32 | 0.00× | 0.0999 | 0.1665 |
| GNG | 10,088 | 155,111 | 404,026 | 107,695.3 ± 6,708.4 | 44.17 | 3.75× | 0.000999 | 0.004701 $\star$ |
| GOR_L | 1,684 | 12,641 | 2,679 | 3,711.6 ± 1,121.3 | -0.92 | 0.72× | 0.3576 | 0.4769 |
| GOR_R | 1,739 | 13,544 | 2,242 | 4,019.3 ± 1,126.5 | -1.58 | 0.56× | 0.1079 | 0.1692 |
| IB_L | 2,772 | 21,250 | 168 | 724.1 ± 292.1 | -1.90 | 0.23× | 0.04396 | 0.08373 |
| IB_R | 2,716 | 22,765 | 169 | 1,037.8 ± 354.6 | -2.45 | 0.16× | 0.02697 | 0.06539 |
| ICL_L | 4,178 | 40,552 | 22,299 | 27,400.1 ± 5,096.8 | -1.00 | 0.81× | 0.3167 | 0.4294 |
| ICL_R | 4,567 | 44,900 | 37,187 | 48,872.9 ± 8,952.0 | -1.31 | 0.76× | 0.1748 | 0.259 |
| IPS_L | 4,043 | 44,537 | 3,328 | 2,999.2 ± 500.1 | 0.66 | 1.11× | 0.4995 | 0.6244 |
| IPS_R | 4,019 | 44,102 | 3,228 | 4,997.1 ± 792.7 | -2.23 | 0.65× | 0.03397 | 0.06967 |
| LAL_L | 2,755 | 34,071 | 18,467 | 26,274.9 ± 3,590.0 | -2.17 | 0.70× | 0.02997 | 0.0685 |
| LAL_R | 2,405 | 31,592 | 14,876 | 36,592.8 ± 4,721.8 | -4.60 | 0.41× | 0.000999 | 0.004701 $\star$ |
| LA_L | 6,570 | 10,482 | 0 | 0.0 ± 0.0 | 0.00 | $\infty \times$ | 1 | 1 |
| LA_R | 8,736 | 17,543 | 0 | 0.0 ± 0.0 | -0.03 | 0.00× | 1 | 1 |
| LH_L | 4,067 | 41,132 | 15,174 | 22,064.7 ± 3,117.2 | -2.21 | 0.69× | 0.02498 | 0.06244 |
| LH_R | 3,977 | 47,102 | 21,262 | 55,690.5 ± 6,924.0 | -4.97 | 0.38× | 0.000999 | 0.004701 $\star$ |

Table 5 continued
| Neuropil | Nodes | Edges | Real | Null (mean ± SD) | Z | Fold | p | q |
| --- | --- | --- | --- | --- | --- | --- | --- | --- |
| LOP_L | 11,211 | 68,368 | 21,894 | 14,181.7 ± 5,604.0 | 1.38 | 1.54× | 0.1039 | 0.1692 |
| LOP_R | 11,796 | 115,374 | 117,313 | 86,262.8 ± 24,163.5 | 1.29 | 1.36× | 0.1828 | 0.2659 |
| LO_L | 22,364 | 219,046 | 740,361 | 181,899.0 ± 44,197.3 | 12.64 | 4.07× | 0.000999 | 0.004701 * |
| LO_R | 22,057 | 232,176 | 780,106 | 188,410.3 ± 42,531.5 | 13.91 | 4.14× | 0.000999 | 0.004701 * |
| MB_CA_L | 3,540 | 20,576 | 135 | 57.9 ± 50.8 | 1.52 | 2.33× | 0.07493 | 0.135 |
| MB_CA_R | 3,426 | 20,118 | 29 | 22.0 ± 25.7 | 0.27 | 1.32× | 0.7712 | 0.8837 |
| MB_ML_L | 4,015 | 23,461 | 39 | 132.9 ± 228.8 | -0.41 | 0.29× | 0.4066 | 0.5246 |
| MB_ML_R | 3,841 | 19,641 | 0 | 59.8 ± 63.9 | -0.94 | 0.00× | 0.1279 | 0.1967 |
| MB_PED_L | 3,805 | 17,489 | 150 | 20.9 ± 26.3 | 4.91 | 7.17× | 0.01099 | 0.0293 * |
| MB_PED_R | 3,954 | 18,605 | 635 | 84.0 ± 43.7 | 12.61 | 7.56× | 0.000999 | 0.004701 * |
| MB_VL_L | 2,765 | 12,957 | 17 | 5.7 ± 17.4 | 0.65 | 2.97× | 0.07592 | 0.135 |
| MB_VL_R | 2,464 | 12,072 | 0 | 2.9 ± 12.6 | -0.23 | 0.00× | 0.8621 | 0.9714 |
| ME_L | 35,624 | 421,151 | 144,012 | 39,902.2 ± 8,097.3 | 12.86 | 3.61× | 0.000999 | 0.004701 * |
| ME_R | 36,142 | 488,839 | 270,338 | 89,732.1 ± 17,267.5 | 10.46 | 3.01× | 0.000999 | 0.004701 * |
| NO | 1,031 | 6,403 | 9 | 668.3 ± 203.9 | -3.23 | 0.01× | 0.006993 | 0.02152 * |
| OCG | 301 | 618 | 0 | 0.0 ± 0.0 | 0.00 | $\infty \times$ | 1 | 1 |
| PB | 1,346 | 5,675 | 4 | 21.7 ± 13.5 | -1.31 | 0.18× | 0.1409 | 0.2126 |
| PLP_L | 7,823 | 73,308 | 27,456 | 73,096.9 ± 14,469.5 | -3.15 | 0.38× | 0.007992 | 0.02283 * |
| PLP_R | 7,383 | 65,621 | 28,920 | 50,801.9 ± 10,360.6 | -2.11 | 0.57× | 0.03297 | 0.06967 |
| PRW | 2,205 | 32,087 | 16,863 | 23,691.0 ± 2,284.4 | -2.99 | 0.71× | 0.006993 | 0.02152 * |
| PVLP_L | 5,446 | 82,315 | 219,208 | 814,846.1 ± 77,567.0 | -7.68 | 0.27× | 0.000999 | 0.004701 * |
| PVLP_R | 4,836 | 71,855 | 115,730 | 496,171.1 ± 53,793.3 | -7.07 | 0.23× | 0.000999 | 0.004701 * |
| SAD | 6,775 | 68,539 | 22,406 | 2,707.6 ± 348.6 | 56.50 | 8.28× | 0.000999 | 0.004701 * |
| SCL_L | 4,955 | 37,251 | 8,791 | 9,516.0 ± 1,868.3 | -0.39 | 0.92× | 0.6933 | 0.8404 |
| SCL_R | 5,542 | 44,354 | 18,944 | 20,101.9 ± 4,169.7 | -0.28 | 0.94× | 0.7702 | 0.8837 |
| SIP_L | 4,399 | 26,299 | 1,123 | 2,336.4 ± 879.6 | -1.38 | 0.48× | 0.1059 | 0.1692 |
| SIP_R | 4,024 | 28,714 | 2,099 | 6,224.8 ± 1,874.7 | -2.20 | 0.34× | 0.03397 | 0.06967 |
| SLP_L | 5,753 | 44,015 | 4,337 | 6,533.3 ± 1,110.1 | -1.98 | 0.66× | 0.03996 | 0.07797 |
| SLP_R | 5,988 | 59,171 | 20,763 | 30,511.6 ± 4,355.9 | -2.24 | 0.68× | 0.02897 | 0.06817 |
| SMP_L | 6,217 | 62,718 | 12,706 | 60,438.6 ± 12,838.2 | -3.72 | 0.21× | 0.006993 | 0.02152 * |
| SMP_R | 6,294 | 69,331 | 21,142 | 77,554.2 ± 12,896.5 | -4.37 | 0.27× | 0.001998 | 0.008413 * |
| SPS_L | 4,917 | 57,941 | 31,159 | 21,384.1 ± 2,622.8 | 3.73 | 1.46× | 0.001998 | 0.008413 * |
| SPS_R | 5,538 | 69,996 | 58,735 | 43,965.9 ± 5,765.0 | 2.56 | 1.34× | 0.02198 | 0.05672 |
| UNASGD | 953 | 713 | 0 | 0.0 ± 0.0 | 0.00 | $\infty \times$ | 1 | 1 |
| VES_L | 2,357 | 27,454 | 18,238 | 12,860.2 ± 1,868.9 | 2.88 | 1.42× | 0.007992 | 0.02283 * |
| VES_R | 2,420 | 29,167 | 16,478 | 19,452.3 ± 2,664.5 | -1.12 | 0.85× | 0.2597 | 0.3583 |
| WED_L | 3,035 | 27,452 | 11,361 | 4,361.8 ± 740.7 | 9.45 | 2.60× | 0.000999 | 0.004701 * |
| WED_R | 3,153 | 31,552 | 9,957 | 2,960.5 ± 504.6 | 13.86 | 3.36× | 0.000999 | 0.004701 * |
| **TOTAL** | **384,417** | **3,869,878** | **6,872,904** | **12,321,112.8 ± 394,661.6** | **-13.80** | **0.56×** | **0.000999** | **—** |

\* = significant at FDR $\alpha$ = 0.05.

**Table 6:** Signed-XOR motif, embedded counting: per-neuropil enrichment in FlyWire *Drosophila, n* = 1000 degree- and Dale-preserving null replicates. ⋆ marks significance at FDR *α* = 0.05. 59/80 neuropils significant.

| Neuropil | Nodes | Edges | Real | Null (mean ± SD) | $Z$ | Fold | $p$ | $q$ |
| --- | --- | --- | --- | --- | --- | --- | --- | --- |
| AL_L | 3,196 | 36,659 | 17,348,520 | 113,177,132.4 ± 25,516,066.3 | -3.76 | 0.15× | 0.003996 | 0.00592 * |
| AL_R | 3,392 | 36,826 | 2,528,501 | 30,044,353.2 ± 8,485,464.6 | -3.24 | 0.08× | 0.006993 | 0.01017 * |
| AME_L | 355 | 1,424 | 6,549 | 88.7 ± 86.4 | 74.76 | 73.87× | 0.000999 | 0.001631 * |
| AME_R | 378 | 1,400 | 12,070 | 731.5 ± 631.5 | 17.95 | 16.50× | 0.000999 | 0.001631 * |
| AMMC_L | 1,376 | 13,398 | 127,329 | 4,843.9 ± 2,175.2 | 56.31 | 26.29× | 0.000999 | 0.001631 * |
| AMMC_R | 1,666 | 12,217 | 7,535 | 90.7 ± 75.2 | 98.94 | 83.04× | 0.000999 | 0.001631 * |
| AOTU_L | 1,456 | 12,139 | 189,097 | 110,734.8 ± 79,561.1 | 0.98 | 1.71× | 0.2208 | 0.2849 |
| AOTU_R | 1,649 | 14,440 | 343,055 | 39,415.5 ± 34,045.5 | 8.92 | 8.70× | 0.000999 | 0.001631 * |
| ATL_L | 1,344 | 6,334 | 244 | 12.5 ± 26.3 | 8.81 | 19.54× | 0.003996 | 0.00592 * |
| ATL_R | 1,284 | 5,838 | 1,532 | 15.6 ± 59.8 | 25.37 | 98.19× | 0.001998 | 0.003134 * |
| AVLP_L | 4,493 | 98,418 | 1,813,119,238 | 130,426,266.7 ± 17,831,250.2 | 94.37 | 13.90× | 0.000999 | 0.001631 * |
| AVLP_R | 5,703 | 131,152 | 2,153,560,129 | 161,438,273.8 ± 22,363,787.3 | 89.08 | 13.34× | 0.000999 | 0.001631 * |
| BU_L | 259 | 632 | 0 | 0.0 ± 0.0 | 0.00 | $\infty \times$ | 1 | 1 |
| BU_R | 350 | 1,043 | 0 | 0.0 ± 0.0 | -0.03 | 0.00× | 1 | 1 |
| CAN_L | 537 | 2,997 | 0 | 1.4 ± 7.2 | -0.20 | 0.00× | 0.8771 | 0.9746 |
| CAN_R | 931 | 3,860 | 0 | 0.0 ± 0.3 | -0.15 | 0.00× | 1 | 1 |
| CRE_L | 5,196 | 36,450 | 20,266,822 | 10,185,633.7 ± 3,370,274.8 | 2.99 | 1.99× | 0.01099 | 0.01542 * |
| CRE_R | 5,067 | 37,359 | 3,436,316 | 7,844,816.2 ± 2,147,778.6 | -2.05 | 0.44× | 0.03397 | 0.04606 * |
| EB | 938 | 17,022 | 67,873,620 | 9,048,737.7 ± 1,504,189.1 | 39.11 | 7.50× | 0.000999 | 0.001631 * |
| EPA_L | 1,629 | 11,651 | 7,097 | 227.5 ± 136.6 | 50.28 | 31.19× | 0.000999 | 0.001631 * |
| EPA_R | 1,401 | 9,605 | 13,090 | 113.3 ± 89.1 | 145.59 | 115.53× | 0.000999 | 0.001631 * |
| FB | 3,127 | 49,145 | 36,065,242 | 2,184,763.0 ± 422,170.2 | 80.25 | 16.51× | 0.000999 | 0.001631 * |
| FLA_L | 1,890 | 14,969 | 36,697 | 77.5 ± 59.3 | 617.19 | 473.67× | 0.000999 | 0.001631 * |
| FLA_R | 2,060 | 17,993 | 37,157 | 408.4 ± 240.7 | 152.67 | 90.97× | 0.000999 | 0.001631 * |
| GA_L | 417 | 1,843 | 4,655 | 45.2 ± 96.1 | 47.99 | 103.00× | 0.000999 | 0.001631 * |
| GA_R | 312 | 1,318 | 7,638 | 897.3 ± 868.9 | 7.76 | 8.51× | 0.000999 | 0.001631 * |
| GNG | 10,088 | 155,111 | 278,928,206 | 41,788.6 ± 6,678.7 | 41,757.81 | 6,674.75× | 0.000999 | 0.001631 * |
| GOR_L | 1,684 | 12,641 | 174,185 | 9,153.2 ± 5,606.3 | 29.44 | 19.03× | 0.000999 | 0.001631 * |
| GOR_R | 1,739 | 13,544 | 258,835 | 11,424.5 ± 6,052.4 | 40.88 | 22.66× | 0.000999 | 0.001631 * |
| IB_L | 2,772 | 21,250 | 1,000 | 260.7 ± 292.4 | 2.53 | 3.84× | 0.02797 | 0.03858 * |
| IB_R | 2,716 | 22,765 | 8,254 | 643.2 ± 804.5 | 9.46 | 12.83× | 0.001998 | 0.003134 * |
| ICL_L | 4,178 | 40,552 | 521,543 | 33,079.5 ± 20,217.8 | 24.16 | 15.77× | 0.000999 | 0.001631 * |
| ICL_R | 4,567 | 44,900 | 2,582,571 | 206,113.6 ± 92,406.9 | 25.72 | 12.53× | 0.000999 | 0.001631 * |
| IPS_L | 4,043 | 44,537 | 833,393 | 1,066.8 ± 470.9 | 1,767.43 | 781.17× | 0.000999 | 0.001631 * |
| IPS_R | 4,019 | 44,102 | 3,573,520 | 2,804.5 ± 1,009.0 | 3,539.03 | 1,274.23× | 0.000999 | 0.001631 * |
| LAL_L | 2,755 | 34,071 | 11,998,959 | 133,430.0 ± 38,588.0 | 307.49 | 89.93× | 0.000999 | 0.001631 * |
| LAL_R | 2,405 | 31,592 | 22,067,445 | 433,852.3 ± 121,734.4 | 177.71 | 50.86× | 0.000999 | 0.001631 * |
| LA_L | 6,570 | 10,482 | 0 | 0.0 ± 0.0 | 0.00 | $\infty \times$ | 1 | 1 |
| LA_R | 8,736 | 17,543 | 0 | 0.0 ± 0.0 | 0.00 | $\infty \times$ | 1 | 1 |
| LH_L | 4,067 | 41,132 | 36,999,593 | 1,368,507.4 ± 431,388.5 | 82.60 | 27.04× | 0.000999 | 0.001631 * |
| LH_R | 3,977 | 47,102 | 25,567,101 | 1,941,941.8 ± 728,743.0 | 32.42 | 13.17× | 0.000999 | 0.001631 * |

Table 6 continued
| Neuropil | Nodes | Edges | Real | Null (mean ± SD) | Z | Fold | p | q |
| --- | --- | --- | --- | --- | --- | --- | --- | --- |
| LOP_L | 11,211 | 68,368 | 1,425,689 | 147,495.4 ± 170,453.2 | 7.50 | 9.67× | 0.002997 | 0.004611 * |
| LOP_R | 11,796 | 115,374 | 139,930,211 | 56,071,403.4 ± 27,064,738.8 | 3.10 | 2.50× | 0.007992 | 0.01142 * |
| LO_L | 22,364 | 219,046 | 7,497,224 | 25,570,973.1 ± 28,767,371.3 | -0.63 | 0.29× | 0.2557 | 0.3148 |
| LO_R | 22,057 | 232,176 | 60,886,602 | 47,713,327.2 ± 27,242,304.9 | 0.48 | 1.28× | 0.5824 | 0.6852 |
| MB_CA_L | 3,540 | 20,576 | 0 | 247.4 ± 334.7 | -0.74 | 0.00× | 0.1309 | 0.1743 |
| MB_CA_R | 3,426 | 20,118 | 2,169 | 4,313.9 ± 3,116.0 | -0.69 | 0.50× | 0.4146 | 0.5025 |
| MB_ML_L | 4,015 | 23,461 | 6,407 | 16,365.8 ± 20,598.6 | -0.48 | 0.39× | 0.4805 | 0.5738 |
| MB_ML_R | 3,841 | 19,641 | 6,118 | 14,889.3 ± 28,846.1 | -0.30 | 0.41× | 0.8012 | 0.9028 |
| MB_PED_L | 3,805 | 17,489 | 2,231 | 17,095.5 ± 31,947.2 | -0.47 | 0.13× | 0.2288 | 0.2905 |
| MB_PED_R | 3,954 | 18,605 | 7,477 | 9,441.1 ± 7,236.1 | -0.27 | 0.79× | 0.6993 | 0.7992 |
| MB_VL_L | 2,765 | 12,957 | 8 | 30,487.5 ± 63,033.4 | -0.48 | 0.00× | 0.1329 | 0.1743 |
| MB_VL_R | 2,464 | 12,072 | 218,876 | 115,932.8 ± 129,843.9 | 0.79 | 1.89× | 0.2418 | 0.3022 |
| ME_L | 35,624 | 421,151 | 2,460,221 | 13,466.2 ± 6,740.3 | 363.00 | 182.70× | 0.000999 | 0.001631 * |
| ME_R | 36,142 | 488,839 | 10,890,500 | 73,907.9 ± 25,065.0 | 431.54 | 147.35× | 0.000999 | 0.001631 * |
| NO | 1,031 | 6,403 | 346,122 | 1,275.5 ± 716.1 | 481.58 | 271.37× | 0.000999 | 0.001631 * |
| OCG | 301 | 618 | 0 | 0.0 ± 0.0 | 0.00 | $\infty \times$ | 1 | 1 |
| PB | 1,346 | 5,675 | 12,986 | 120.7 ± 166.4 | 77.31 | 107.62× | 0.000999 | 0.001631 * |
| PLP_L | 7,823 | 73,308 | 561,372 | 57,947.3 ± 22,349.9 | 22.52 | 9.69× | 0.000999 | 0.001631 * |
| PLP_R | 7,383 | 65,621 | 864,896 | 39,255.8 ± 16,249.9 | 50.81 | 22.03× | 0.000999 | 0.001631 * |
| PRW | 2,205 | 32,087 | 6,974,235 | 137,195.1 ± 30,446.3 | 224.56 | 50.83× | 0.000999 | 0.001631 * |
| PVLP_L | 5,446 | 82,315 | 105,691,616 | 14,660,065.3 ± 3,589,253.1 | 25.36 | 7.21× | 0.000999 | 0.001631 * |
| PVLP_R | 4,836 | 71,855 | 60,808,493 | 5,226,053.2 ± 1,628,857.1 | 34.12 | 11.64× | 0.000999 | 0.001631 * |
| SAD | 6,775 | 68,539 | 1,532,400 | 279.6 ± 113.5 | 13,504.69 | 5,481.47× | 0.000999 | 0.001631 * |
| SCL_L | 4,955 | 37,251 | 398,992 | 16,490.4 ± 7,231.7 | 52.89 | 24.20× | 0.000999 | 0.001631 * |
| SCL_R | 5,542 | 44,354 | 364,434 | 46,744.8 ± 20,331.7 | 15.63 | 7.80× | 0.000999 | 0.001631 * |
| SIP_L | 4,399 | 26,299 | 467,368 | 606,844.6 ± 305,792.6 | -0.46 | 0.77× | 0.6074 | 0.7042 |
| SIP_R | 4,024 | 28,714 | 2,889,595 | 60,189.5 ± 34,232.1 | 82.65 | 48.01× | 0.000999 | 0.001631 * |
| SLP_L | 5,753 | 44,015 | 562,823 | 9,518.2 ± 3,896.4 | 142.00 | 59.13× | 0.000999 | 0.001631 * |
| SLP_R | 5,988 | 59,171 | 4,146,074 | 94,513.3 ± 31,497.2 | 128.63 | 43.87× | 0.000999 | 0.001631 * |
| SMP_L | 6,217 | 62,718 | 21,568,387 | 3,005,375.7 ± 1,301,547.8 | 14.26 | 7.18× | 0.000999 | 0.001631 * |
| SMP_R | 6,294 | 69,331 | 14,930,901 | 2,867,480.9 ± 1,023,766.3 | 11.78 | 5.21× | 0.000999 | 0.001631 * |
| SPS_L | 4,917 | 57,941 | 1,309,351 | 17,096.3 ± 5,516.1 | 234.27 | 76.59× | 0.000999 | 0.001631 * |
| SPS_R | 5,538 | 69,996 | 3,746,567 | 27,682.0 ± 8,186.2 | 454.29 | 135.34× | 0.000999 | 0.001631 * |
| UNASGD | 953 | 713 | 0 | 0.0 ± 0.0 | 0.00 | $\infty \times$ | 1 | 1 |
| VES_L | 2,357 | 27,454 | 2,041,430 | 29,474.0 ± 10,664.5 | 188.66 | 69.26× | 0.000999 | 0.001631 * |
| VES_R | 2,420 | 29,167 | 2,644,902 | 34,500.1 ± 12,101.5 | 215.71 | 76.66× | 0.000999 | 0.001631 * |
| WED_L | 3,035 | 27,452 | 6,112,571 | 6,123.4 ± 2,767.8 | 2,206.25 | 998.24× | 0.000999 | 0.001631 * |
| WED_R | 3,153 | 31,552 | 6,009,967 | 3,483.8 ± 1,558.7 | 3,853.46 | 1,725.11× | 0.000999 | 0.001631 * |
| **TOTAL** | **384,417** | **3,869,878** | **4,965,825,953** | **625,438,802.7 ± 63,073,546.1** | **68.81** | **7.94×** | **0.000999** | **—** |

\* = significant at FDR $\alpha = 0.05$ .

### 4.4 Mouse V1: layer-structured over-representation with a clean negative control

In a biophysically detailed model of mouse V1 [15] (230,922 neurons, ~ 7 × 10^7^ synapses), the signed-XOR motif counted locally around 60 inhibitory pivots (~5 per cell type, *N* = 100 nulls each; Section C, Table 7) is strongly over-represented in layers 2/3, 4 and 5: per-pivot median folds of 15–852×, single-pivot *Z* up to +7616, and a global enrichment of 315× (32.4M real motifs vs. 102,712 expected across the 60 subgraphs). Layer 4 leads, two orders of magnitude above *Drosophila* AVLP_L, and the motif is conspicuously *absent* in layer 6 (all three inhibitory types non-significant: per-pivot |*Z*| < 2, *p* ≥ 0.13, real counts of 0–357). The same counter applied uniformly across all twelve cell types switches off cleanly at the L5/L6 boundary, where subgraph sizes are comparable, making the layer-6 null a strong internal control against a combinatorial artefact.

**Table 7.** Signed-XOR over-representation in the Allen mouse V1 model, aggregated by inhibitory cell type over 60 pivot neurons (~5 per type) at |*w*| ≥ 0.1, *k* = 2, *N* = 100 null replicates per pivot. “Real” and “Null mean” are summed/averaged over the pivots of each type; “Fold” and “*Z*” are per-pivot medians, “*Z*_max_” the largest single-pivot value; “*p*” is the median empirical value (floor 1*/*101 ≈ 0.0099). Bold = significant; layer-6 rows are non-significant. GLOBAL is the pooled real/null count over all 60 subgraphs.

| Layer | Cell type | Real | Null mean | Fold | Z | $Z_{\max}$ | p |
| --- | --- | --- | --- | --- | --- | --- | --- |
| L2/3 | i23Pvalb | 1,039,750 | 13,012 | **104.0**× | +523 | +782 | 0.0099 |
| L2/3 | i23Sst | 494,303 | 5,265 | **177.3**× | +677 | +1018 | 0.0099 |
| L2/3 | i23Htr3a | 121,209 | 280 | **375.6**× | +502 | +1719 | 0.0099 |
| L4 | i4Pvalb | 11,122,740 | 20,100 | **515.9**× | +2663 | +7616 | 0.0099 |
| L4 | i4Sst | 9,821,057 | 40,208 | **208.8**× | +1196 | +2219 | 0.0099 |
| L4 | i4Htr3a | 9,164,822 | 17,269 | **852.0**× | +3780 | +6291 | 0.0099 |
| L5 | i5Pvalb | 20,550 | 2,185 | **14.9**× | +47 | +57 | 0.0099 |
| L5 | i5Sst | 608,700 | 3,605 | **174.8**× | +577 | +963 | 0.0099 |
| L5 | i5Htr3a | 2,910 | 92 | **41.8**× | +40 | +100 | 0.0099 |
| L6 | i6Pvalb | 357 | 655 | 0.4× | −1.0 | +1.4 | 0.13 (n.s.) |
| L6 | i6Sst | 4 | 22 | 0.0× | −0.4 | −0.0 | 0.83 (n.s.) |
| L6 | i6Htr3a | 10 | 20 | 0.2× | −0.4 | +0.1 | 0.73 (n.s.) |
| **GLOBAL** | | **32,396,412** | **102,712** | **315**× | — | +7616 | — |

**Table 8:**
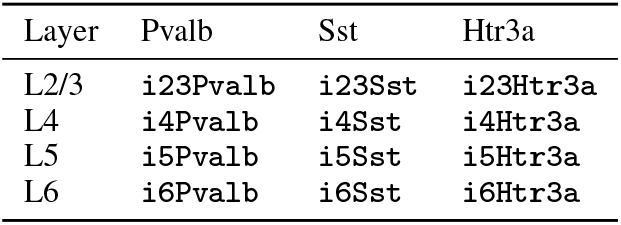
Inhibitory cell types used as role-6 (INH hub) candidates in the mouse V1 motif search.

The same layer structure is found in the axon-proofread subset of the EM-reconstructed MICrONS connectome [16] (1,708 neurons, 86,793 signed edges). Here layer 4 again dominates (46% of motifs, led by `i4Pvalb` with 55,870 motifs), layers 2/3 and 5 are intermediate (28% and 25%), and layer 6 contributes only 1.4% of motifs despite a fully populated complement of layer-6 interneurons (Section C, Table 9). We restrict this analysis to the proofread subset, in which each source neuron’s out-degree is real by construction, rather than imposing an out-degree ceiling on the full graph: a real layer-4 parvalbumin interneuron makes of order several thousand output synapses and hence several hundred distinct targets, so a low ceiling removes genuine connectivity rather than merge artefacts. Unlike the Allen model, we make no enrichment claim for MICrONS: on this small, dense, spatially organised graph a degree- and cell-type-preserving null produces *more* signed-XOR motifs than the real graph (of order 6 × 10^5^ versus 1.6 × 10^5^), so it does not serve as a chance baseline (Section C). Notably, the intermediate-layer ordering differs between substrates: L2/3 exceeds L5 in the Allen model, whereas L5 exceeds L2/3 in the MICrONS connectome, and layer 5 is moreover the only layer in which the Sst pivot exceeds the Pvalb pivot.

**Table 9.** Signed-XOR motif counts in the axon-proofread MICrONS EM-reconstructed mouse V1 connectome (1,708 neurons, 86,793 signed edges; 10 inhibitory pivots sampled per cell type), by inhibitory cell-type group and cortical layer, with layer assignment taken from the MICrONS cell-type labels. Layer 4 dominates and is led by i4Pvalb; layer 5 is the only layer in which the Sst pivot exceeds the Pvalb pivot; layer 6 contributes 1.4% of motifs despite a fully populated layer-6 interneuron complement. We do not report a null mean here: on this small, dense, spatially organised graph a degree- and cell-type-preserving null produces more motifs than the real graph (see text), so it does not serve as a chance baseline; a distance-preserving null is identified as follow-up work.

| Layer | Cell-type group | Real motifs |
| --- | --- | --- |
| L2/3 | i23* (all) | 44,060 |
| L4 | i4Pvalb | 55,870 |
| L4 | i4Sst | 16,830 |
| L4 | i4Htr3a | 1,231 |
| L5 | i5Pvalb | 15,467 |
| L5 | i5Sst | 23,369 |
| L5 | i5Htr3a | 393 |
| L6 | i6* (all) | 2,247 |
| Total |  | 159,467 |

### 4.5 Cross-species synthesis

The same eight-node, twelve-edge topology, defined purely as a connectivity template with no biological detail beyond edge sign, is significantly over-represented as an embedded subgraph in all three networks, which span four orders of magnitude in size (3,186 signed edges in the worm; ~10^5^ in an AVLP fly neuropil; ~4 × 10^7^ in thresholded mouse V1) and three different dominant-neurotransmitter regimes (a glutamate-driven worm, an acetylcholine-driven fly, a glutamate–GABA cortical model). Two further observations qualify this robustness. First, the enrichment is anatomically structured rather than uniform: in mouse V1 it is strong in the intracortical processing layers (2/3–5) and absent in the corticothalamic feedback layer (6), a layer-resolved pattern also found in the axon-proofread EM-reconstructed MICrONS connectome (layer-4-dominant, parvalbumin-led, layer 6 strongly under-represented), that is, in an experimentally measured rather than simulated network; we note that the MICrONS pattern is reported as layer structure of the motif and not as enrichment over a null, since a degree-preserving null is not an adequate chance model on that dense, spatially embedded graph (Section C); in *Drosophila* it is strongest in the gnathal/saddle and inferior-posterior regions and reversed (depleted) in the antennal lobes. Second, the specificity for the *signed* extension over the bare XOR core is itself species-dependent: in the fly the embedded signed-XOR is ~3.3× more enriched than the bare core (Table 4), i.e. the feedback module is the discriminating feature, whereas in the worm both the bare and signed motifs are strongly enriched, with the bare core even more so.

We make no causal claim that the motifs detected here are functionally engaged in any specific computation; that would require interventional experiments beyond connectome-level structural analysis. What these results establish is a scale-spanning, anatomically structured topological bias in favour of the signed-XOR template under the null models tested here, with implications for the inductive biases that artificial architectures for local credit assignment might recapitulate.

## 5 Circuit-level functional validation in a spiking model

The enumeration results of Section 4 establish that the signed-XOR *topology* is over-represented across three connectomes, and that in mouse V1 (Section 4.4) this over-representation is layer- and cell-type-structured. Structural enrichment, however, does not establish that the topology actually computes a signed error under realistic dynamics: as noted in the Limitations, this requires spiking simulation with biological time constants. This section supplies that test for the *isolated* eight-neuron motif. The goal is deliberately narrow: not to simulate V1, but to verify that the circuit of Section 2 performs the signed-XOR computation when instantiated as spiking neurons, and to ask which inhibitory cell types can dynamically support it. The latter question turns out to connect directly to the layer-specific cell-type structure of Section 4.

### 5.1 Model

We instantiate the eight roles {*I, P, Li, Lp*, INH, XOR, *F* ^+^, *F* ^−^} of Section 2 as one current-based leaky integrate-and-fire (LIF) neuron each, in Brian2 [18], wired with the twelve directed edges of the motif. Two specific design commitments make the test concrete:

- **Variant B (intermediate-side feedback)**. The two error aggregators read the sign from the XOR-internal relays: *Li* → *F* ^+^ and *Lp* → *F* ^−^, together with the shared edges XOR → *F* ^+^ and XOR → *F* ^−^.
- **Feedforward inhibition**. The pivot inhibits the relays (INH ⊣ *Li*, INH ⊣ *Lp*), not the inputs, and is tuned as a *coincidence detector*: a single active input is sub-threshold at the pivot, and only simultaneous input drive makes it fire and veto the relays. This is the feedforward perisomatic-inhibition regime that distinguishes the motif from recurrent winner-take-all (see the discussion of lateral inhibition and winner-take-all below), and the regime in which fast-spiking inhibition enforces a narrow coincidence window [19, 20].

Membrane time constants are assigned per Allen V1 cell type: thalamo-recipient inputs as `e4other` (*τ*_*m*_ = 12 ms), relays and comparator as `e23Cux2` (*τ*_*m*_ = 15 ms), and the inhibitory pivot as a fast-spiking `i4Pvalb` interneuron (*τ*_*m*_ = 6 ms, refractory 1 ms). Each trial drives the active input(s) for 100 ms and a role is scored “active” at ≥ 2 spikes. The simulation uses Brian2’s pure-NumPy code-generation target, requires no compiler, and runs in seconds on a laptop; full parameters, weights and the integration scheme are in the released code (Section 7).

The motif’s twelve synapses fall into two classes. Six are **drive edges** (*I* → *Li, P* → *Lp*, INH ⊣ *Li*, INH ⊣ *Lp, Li* → XOR, *Lp* → XOR): in this simplified model a single presynaptic spike suffices to fire the postsynaptic neuron. The remaining six are **coincidence edges** targeting the three coincidence detectors INH, *F* ^+^ and *F* ^−^, each of which requires both its inputs to arrive together. The circuit therefore is modeled with only two synaptic weights, *W*_*d*_ for drive edges and *W*_*c*_ for coincidence edges, and its behaviour depends only on their ratio. We use *W*_*d*_*/W*_*c*_ = 3 (*W*_*d*_ = 84, *W*_*c*_ = 28 in normalised LIF units), the operating point at which the truth table is reproduced robustly (Section 5.2); the functional window of this ratio is discussed below.

### 5.2 The circuit computes the signed-XOR truth table

Driving the two inputs through all four binary conditions reproduces the signed-XOR truth table deterministically (Table 2, Figure 2). When the inputs agree the circuit rests in homeostasis: the silent case (0, 0) produces no spikes, and in the matched-active case (1, 1) the pivot fires (8 spikes), vetoes both relays, and the comparator and both error neurons stay silent. When the inputs disagree the comparator fires (4 spikes in both mismatch cases) and exactly one error channel responds: the under-shoot case (1, 0) activates *F* ^+^ (3 spikes, *F* ^−^ silent), the driver of potentiation, while the over-shoot case (0, 1) activates *F* ^−^ (7 spikes, *F* ^+^ silent), the driver of depression. The directional decomposition the motif was designed to provide (Section 2) thus holds at the level of individual spikes, not only in the rate abstraction. Under a 20 Hz Poisson background on the input neurons (25 random seeds) the complete four-row table is recovered on 88% of trials, confirming the computation is robust to stochastic drive.

**Figure 2.**
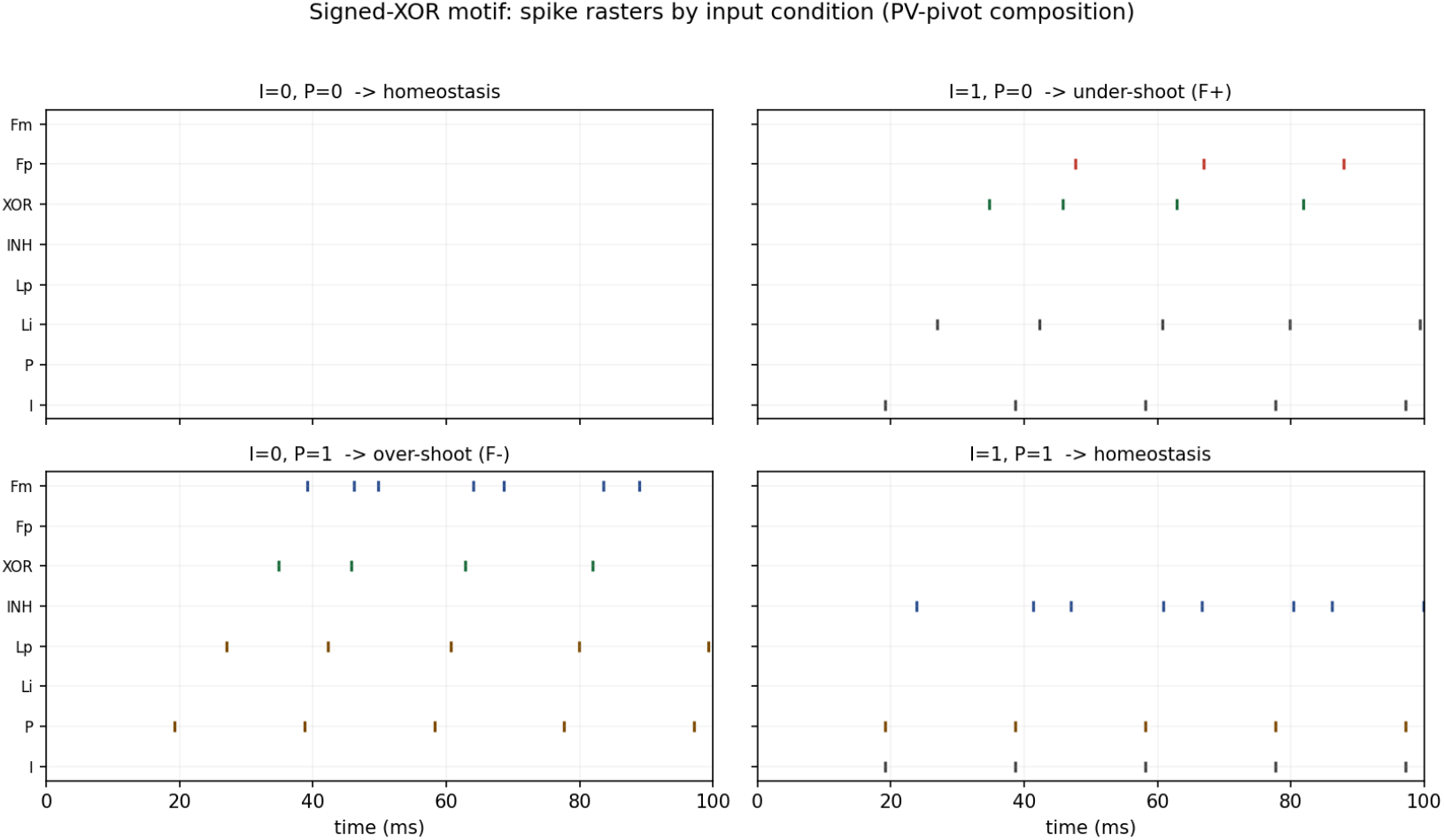
Spike rasters of the eight motif neurons for the four input conditions. The pivot (INH) fires only when both inputs are co-active (1, 1) and silences the relays; the comparator (XOR) fires only on mismatch; *F* ^+^ and *F*^−^ each respond to exactly one mismatch direction.

### 5.3 The feedforward comparator requires a fast-spiking pivot

The coincidence/veto computation depends on the pivot’s membrane integration window being short relative to the synaptic time scale: a slow pivot integrates a *single* input to threshold and fires spuriously, collapsing the exclusive-or. We tested this directly by sweeping the pivot’s membrane time constant 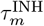 from 4 to 15 ms while holding all else fixed, scoring the full truth table over 15 noisy seeds per value (Figure 3). The circuit succeeds on 100% of trials for 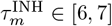 ms (fast-spiking, PV-like), degrades at 8 ms (60%), and fails completely (0%) for 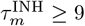 ms, i.e. at the membrane time constants characteristic of pyramidal cells and of the slower Sst and Htr3a/VIP interneuron classes. Consistently, instantiating the pivot as `i4Pvalb` yields a working circuit, whereas instantiating it with an `i4Sst`-like slow pivot at otherwise identical parameters does not.

**Figure 3.**
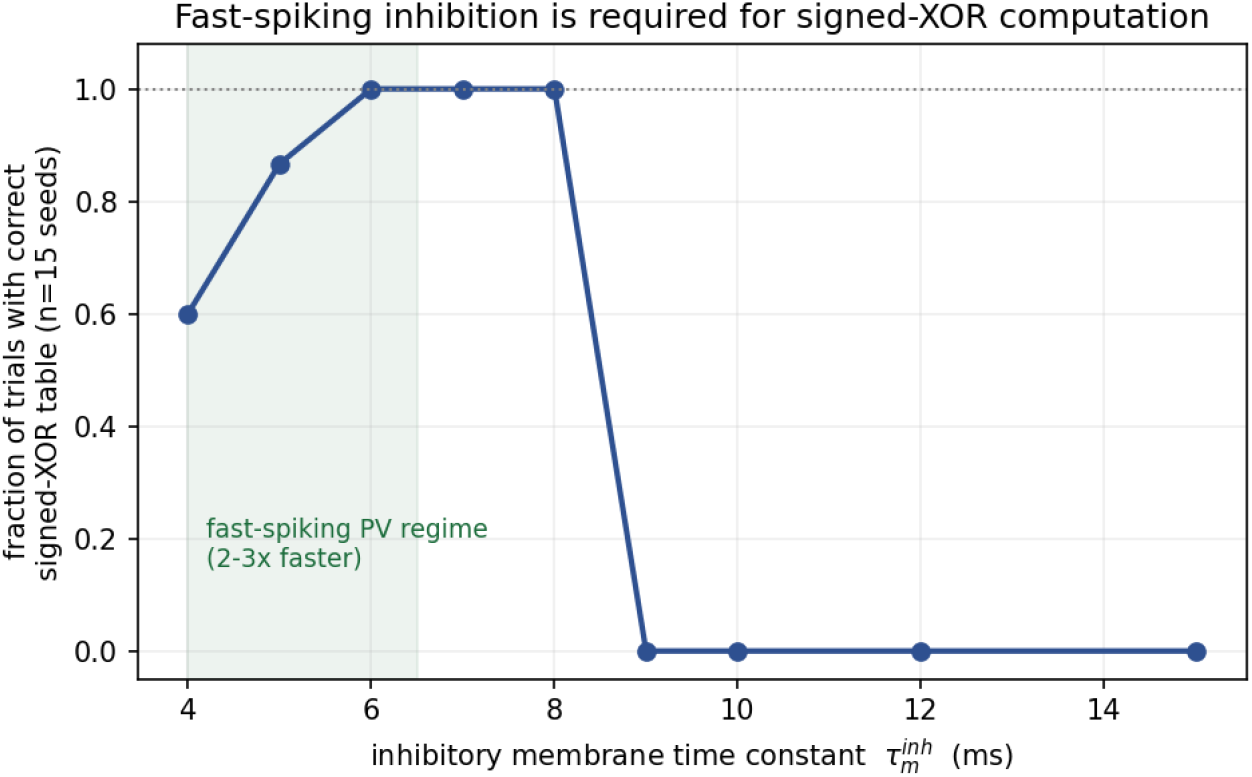
Truth-table success rate (15 noisy seeds per point) versus the pivot’s membrane time constant. The functional window (100%) is confined to the fast-spiking, parvalbumin-like regime (*τ*_*m*_ ≤ 7 ms); the feedforward comparator collapses at the slower time constants of Sst, Htr3a/VIP and pyramidal cells.

This dynamical constraint aligns with the layer- and cell-type structure of the mouse-V1 enrichment (Section 4). There, a strong *Pvalb*-pivoted signal is found in layer 4 (`i4Pvalb`), the canonical site of fast feedforward perisomatic inhibition and lateral suppression [19, 20]; layer 2/3 is instead dominated by Htr3a/VIP and layer 5 by Sst/Htr3a. Our simulation shows that the *feedforward coincidence-comparator* realisation of the signed-XOR, the version implemented here with the pivot directly vetoing the relays, is dynamically available only to fast-spiking pivots.

Among the structurally enriched signed-XOR instances, those that implement the motif as a *fast feedforward comparator* (pivot directly gating the relays within a single gamma cycle) require a fast-spiking parvalbumin pivot and are therefore concentrated in the L4 `i4Pvalb` population. The L2/3 (Htr3a/VIP-dominated) and L5 (Sst-dominated) enrichments cannot operate by this mechanism at their interneurons’ membrane time constants; if functional, they must realise the signed comparison by a different dynamical route, VIP→Sst *disinhibition* in L2/3 [21, 22] or slow dendritic

Sst-mediated inhibition in L5, which the present feedforward model does not capture. The prediction is testable both *in silico* (a cell-type-resolved spiking simulation of the Allen V1 model should find fast coincidence-style mismatch signalling specifically on L4 PV-pivoted instances) and *in vivo* (optogenetic silencing of PV versus Sst/VIP interneurons should abolish mismatch responses in a layer-specific manner).

This sharpens, rather than weakens, the structural finding: the cell-type heterogeneity across layers reported in Section 4 is exactly what one expects if a single connectivity template is being read out by different dynamical mechanisms in different layers, with the fast feedforward mechanism validated here confined to the PV-rich L4.

### 5.4 The error signal is graded and correctly signed

Finally we tested the graded regime. Holding the “target” input *I* fixed and sweeping the “prediction” *P* from 0 to 1.6× its level, *F* ^+^ is active across the entire under-shoot side (prediction below target) and falls silent once the prediction reaches the target, while *F* ^−^ is silent on the under-shoot side and activates on the over-shoot side (Figure 4). The near-silent cross-over occurs at ≈0.9× the target, recovering the homeostatic stopping condition at match. The two channels therefore encode the sign of the mismatch with the correct polarity, as required by the directional-feedback role of *F* ^+^*/F* ^−^ (Section 2).

**Figure 4.**
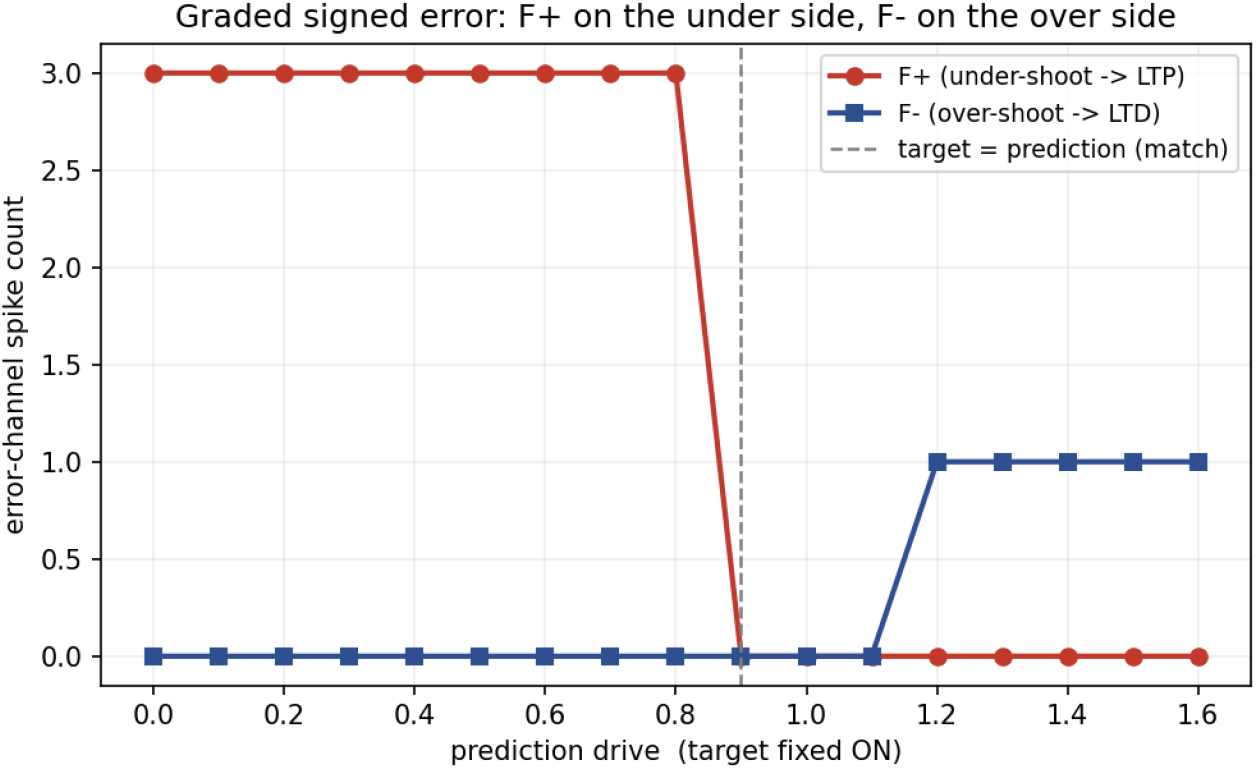
Graded signed error. With the target fixed, *F* ^+^ (potentiation driver) is active when the prediction under-shoots the target and *F*^−^ (depression driver) when it over-shoots, with a near-silent cross-over at match. The motif reads out the sign of the prediction error without any global computation.

### 5.5 Scope of the spiking validation

This is a functional validation of the *isolated* eight-neuron motif, not of V1. It uses one neuron per role, current-based LIF dynamics with cell-type membrane time constants, hand-set synaptic weights, and the Variant B, feedforward-inhibition realisation. It deliberately does *not* include spike-timing-dependent plasticity, dendritic nonlinearities, conductance-based synapses, short-term depression, the alternative disinhibitory (VIP → Sst) and slow-dendritic mechanisms implicated for L2/3 and L5 above, or the embedding of the motif in the full recurrent V1 graph. It establishes that the signed-XOR computation is *realisable* in spiking dynamics with biologically plausible time constants, and that the feedforward realisation is contingent on fast-spiking inhibition; it does not establish that any particular V1 instance is used this way *in vivo*. Coupling the validated motif to the Allen V1 reservoir with cell-type-specific rates and STDP, and modelling the L2/3/L5 non-PV mechanisms, are the natural next steps (Section 6, future work).

## 6 Discussion

### 6.1 The connectomic signature of directional feedback

The central empirical result is not merely that the signed-XOR motif exists in biological connectomes, but that it appears more often than expected under null models that preserve local degree and sign structure. The cleanest within-dataset comparison is the embedded 2 × 2 analysis of *Drosophila* AVLP_L: the signed motif is enriched 13.9×, whereas the bare XOR core is enriched 4.2×. This does not prove that the motif is used for learning, but it indicates that the additional feedback channels are not a trivial consequence of the XOR core alone. In the context of our autoencoder rule [10], this distinction is important: the bare XOR supplies error magnitude, while the signed-XOR supplies a local representation of update direction using two Dale-compatible channels.

### 6.2 Homeostatic error cancellation

The signed-XOR motif also gives a circuit interpretation to homeostasis. In matched states, activity is not simply passed forward: it is cancelled by the inhibitory pivot before reaching the mismatch readout. In mismatched states, cancellation is incomplete and a directional error is exported. This is a local version of a broader predictive-processing principle: predictable activity should be suppressed, whereas prediction errors should remain available for learning. The proposed motif should therefore be understood less as an isolated logical gate and more as a small excitation–inhibition module that prevents redundant amplification while preserving informative deviations.

### 6.3 Why induced counts are depleted: a methodological note

Induced subgraph counts are depleted relative to the null in dense connectomes (Tables 4 and 5). The induced count requires the absence of all extra edges among the motif’s nodes; in a real connectome, motif-participating neurons are densely interconnected with additional edges that disqualify the strict count, whereas randomised graphs have fewer such “almost-motifs”. Induced counting therefore systematically under-represents motif prevalence in dense networks. We report embedded counts as the primary measure and induced counts only as a complementary, conservative check.

### 6.4 Variants A and B: two designs for the same signed-error motif

The motif’s two variants differ in where the sign-defining feedback is read out: Variant A from the inputs, Variant B from the intermediate convergent neurons. Both produce identical logical behaviour. Empirically discriminating between them requires the per-variant counts in real and null graphs (the counter records A/B/AB); a strongly skewed A:B ratio would indicate one is biologically preferred, roughly equal frequencies that they are alternative implementations. *(This per-variant breakdown is work in progress*.)

### 6.5 Distinction from lateral inhibition and Winner-Take-All

Two canonical inhibitory motifs, lateral inhibition and Winner-Take-All (WTA), share the signed-XOR’s lateral-inhibition substrate, raising the question of whether we have merely rediscovered one of them. We argue not. *Lateral inhibition* [23] sharpens relative activity but neither selects among competitors nor exports a categorical signal. *WTA* [24, 25] computes an argmax read out at the competitors themselves (the surviving unit’s activity *is* the answer). The signed-XOR reuses the lateral-inhibition substrate but differs in three ways: the inhibition is *feedforward* (node 6 inhibits the relays 2/4, not the inputs 1/3, so competition resolves one synapse downstream); the readout is *shared* (the integrator, node 5, sits below both lanes and yields a single binary mismatch rather than per-lane survival); and the result is *exported* (the F^+^/F^−^ aggregators broadcast the winning lane as a signed message upstream). Informally, WTA broadcasts “who won” implicitly by silencing losers in place; the signed-XOR broadcasts it explicitly as a labelled message to a different population. This also clarifies why XOR, not argmax, is the right lens: a single linear unit can implement two-input WTA but probably cannot implement two-input XOR [26], and the INH node is formally the hidden unit lifting the XOR node into the non-linearly-separable regime.

### 6.6 Relation to STDP and dendritic XOR computation

An XOR-like operation can be realised at three levels of organisation, which are complementary rather than competing. *STDP* [27–29] is a synaptic learning rule, not a computation; in our framework it is a candidate substrate-builder that could shape the relevant wiring during development, but does not itself produce the eight feedforward edges of the XOR core. *Dendritic XOR* places the computation inside a single neuron [30–33], at zero synaptic-delay cost but under a strict anatomical constraint: the inputs must physically converge on the same neuron. The signed-XOR motif removes this constraint (its input neurons may be anywhere) at the cost of at least eight neurons and twelve synapses. What the motif uniquely provides, and what no synaptic rule or single-neuron dendritic computation supplies on its own, is a *sign-decoded, lane-labelled, exportable* report: a single axon cannot, by definition, broadcast two opposite-sign signals to non-overlapping targets, whereas the F^+^/F^−^ split does exactly that. We therefore read the three mechanisms as a layered hierarchy (STDP shapes wiring; dendritic nonlinearities sharpen local gating; the signed-XOR circuit assembles directional feedback that can be communicated across regions), of which an enumeration of static directed signed subgraphs can detect only the third.

### 6.7 Connections to existing learning frameworks

The motif is compatible with several biologically motivated learning frameworks. *Predictive coding* [5] computes signed prediction errors as paired prediction/reconstruction activity; the signed-XOR is one explicit single-neuron-resolution construction. *Burst-mediated plasticity* [6] conveys teaching signals whose sign the F^+^/F^−^ burst statistics could carry. *Random feedback alignment* [4], used in Peña Fernández et al. [10], needs a fixed random projection of the signed error, which long-range axons from F^+^/F^−^ could provide.

### 6.8 Limitations

#### Sign assignment

The neurotransmitter-based sign rules for *Drosophila* are convenient defaults, not absolute; a receptor-grounded assignment would be a useful extension. The *C. elegans* analysis already uses curated per-edge signs (Section A).

#### Null model

Roberts–Coolen randomisation preserves per-sign degree exactly, but not (in the cross-invertebrate analyses) cell type, distance, spatial embedding, laminar organisation, synaptic weight distributions, or lower-order motif structure. For the MICrONS analysis we additionally layered the null by mapped cell type; this remains a degree-preserving null that does not control distance, laminar position or weight distribution, and on the dense, spatially embedded proofread MICrONS graph it produces more motifs than the real network, so we report the MICrONS result as a description of real, layer-structured connectivity rather than as enrichment over chance (Section C). A distance- and laminar-preserving null, which would provide the appropriate chance baseline for a spatially embedded cortical connectome, is the priority follow-up. For the invertebrate and Allen-model substrates, the reported fold enrichments should be interpreted as enrichment relative to the specific degree- and sign-preserving null used here.

#### Structure versus function

Structural over-representation does not establish functional use. The Brian2 model shows that the isolated motif can implement signed-XOR dynamics under biologically plausible fast-spiking inhibition, and previous autoencoder simulations show that the signed-XOR rule can support learning. Neither result demonstrates that any individual motif instance is used for credit assignment in vivo. That bridge will require cell-type-resolved activity, perturbation and plasticity experiments.

In fact, a ratio sweep at fixed coincidence synaptic weights *W*_*c*_, reveals that the motif is functional only within a narrow window, *W*_*d*_*/W*_*c*_ ∈ [2.75, 3]; outside this window the truth table either fails because coincidence detectors do not reach threshold or because the drive overwhelms the homeostatic veto at match. The narrowness of this band would imply that only motif instances whose drive synapses are roughly three times stronger than their coincidence synapses can support the signed-XOR computation; however this applies only to motifs in isolation, and not highly embedded in a network, where the activation requirements are likely totally different.

### 6.9 On-going work

In parallel to this work, our group is extending the computational results obtained using the signed XOR as a loss function in a shallow autoencoder from the processing of static images like MNIST, to the prediction in temporal sequences like songs and piano music (in preparation); and it is also exploring the dynamic implications for the structuring of memory (in preparation). Our final goal is to test the feasibility of a large neuromorphic computational neural network re-using the mouse connectome, guided by the hypothesised role of the signed XOR motif to generate local credit-assignment signals during learning (in preparation).

## 7 Conclusion

We introduced the signed-XOR motif as a candidate circuit primitive for local homeostatic error cancellation and directional error signalling. The motif extends an XOR mismatch detector with two Dale-compatible feedback channels that distinguish potentiating from depressing updates. Across *Drosophila, C. elegans* and mouse V1 datasets, the motif is enriched relative to sign- and degree-preserving null models, and its distribution is anatomically structured rather than uniform. A spiking Brian2 implementation further shows that the isolated circuit can compute the signed-XOR truth table, produce a graded signed error, and operate in a fast-spiking parvalbumin-like regime. These results do not prove functional deployment in vivo, but they provide a concrete, testable connectomic candidate for local, directional error signalling.

## Data and Code Availability

The code implementing the three signed-XOR motif counters (Allen/MICrONS, FlyWire and *C. elegans* edge-list formats), the Roberts–Coolen randomiser (including the *C. elegans* variant that preserves curated edge signs), and the Brian2 spiking model of a single signed-XOR motif with its functional-validation tests and result figures, with per-component documentation, are available at https://github.com/jesusmarcodelucas/signed-xor-motifs [12]. No connectome data is redistributed; each dataset must be obtained from its original provider under the corresponding terms (see repository documentation).

## Acknowledgments

We thank the Worms, FlyWire and MICrONS consortia for making their connectomic resources available to the community.

## A *C. elegans*: data, methods and results

### Connectome and edge classification

We use the hermaphrodite connectome of Cook et al. [13]. After removing self-loops, the network comprises 300 neurons and 3,669 directed chemical synaptic connections: 2,933 excitatory, 253 inhibitory, and 483 aminergic/unclassified connections that do not enter the motif and were excluded. A key difference from *Drosophila* is the assignment of sign. In the worm, glutamate is predominantly *excitatory*; moreover a subset of aminergic synapses (serotonergic, tyraminergic, octopaminergic) are curated as excitatory, while a minority of cholinergic synapses are curated as unclassified. The neurotransmitter therefore does not determine the functional sign edge-for-edge. We consequently take the excitatory/inhibitory label for every edge *directly from the curated per-edge classification column*, not from neurotransmitter, ensuring that the curated sign is used identically in the empirical network and in every null replicate (Section D).

### Counting mode

Because the connectome is small and densely clustered, we count non-induced (embedded) occurrences: on a graph this dense the induced criterion rejects essentially all empirical instances (the real induced count is zero), and is uninformative.

### Results

Table 3 reports both motifs at *n* = 1000. The bare XOR motif occurs 47,397 times against a null of 1,217.5 ± 64.3 (38.9×, *Z* = 126.8). The signed-XOR motif occurs 110,399 times against 4,536.7 ± 2,027.6 (24.3×, *Z* = 52.2). Both empirical *p* are at the floor 1/1001. Unlike the fly, the nematode strongly favours XOR-type connectivity even without the signed-feedback completion.

## B *Drosophila*: data, methods and full per-neuropil results

### Data and edge sign

We use the proofread FlyWire connectome [14] with neurotransmitter predictions from Eckstein et al. [11]. Edges are classified excitatory if the predicted neurotransmitter is `ACH` and inhibitory if `GABA` or `GLUT` (glutamate is inhibitory in *Drosophila* via GluCl receptors); `SER`, `DA`, `OCT` and others are tracked but excluded from the motif’s required edges and matter only for the strict induced check. Each neuropil is analysed separately using the spatial annotations of Dorkenwald et al. [14]; the null randomisation is layered per neurotransmitter (Section D).

### Validation

The C counter was validated against an independent NetworkX pipeline on AVLP_L: both report exactly 2,098,224 raw mappings (= 1,049,112 distinct after the 2-fold XOR symmetry) for the bare XOR under the induced criterion, agreeing to the unit.

### AVLP_L reference matrix

Table 4 gives the full 2 × 2 of motif type by counting mode at *n* = 1000 on AVLP_L (*N* = 4,493 neurons; *E* = 98,418 synapses: 60,356 excitatory, 36,246 inhibitory, 1,816 other). In embedded mode both motifs are enriched and the signed motif (13.9–14.0×) exceeds the bare core (4.2×); in induced mode both are depleted (the dense-network signature, Section 6.3).

### Full per-neuropil scans

Table 5 (bare XOR, induced) and Table 6 (signed-XOR, embedded) give all 80 neuropils at *n* = 1000 with Benjamini–Hochberg FDR correction. The bare-XOR induced scan is significant in 30*/*80 neuropils (aggregate 0.56×, the induced depletion signature); the signed-XOR embedded scan is significant in 59*/*80 (aggregate 7.94×), with the strongest enrichments in GNG, SAD, the inferior-posterior slope (IPS) and wedge (WED), and significant depletion confined to the antennal lobes and CRE_R.

### Regional interpretation

The embedded signed-XOR enrichment is not uniform. Four broad regimes emerge: most central regions (e.g. SMP, SIP, ICL, LAL) are XOR-neutral-to-depleted under induced counting yet strongly signed-enriched; the lobula (LO_L/R) and mushroom-body peduncle show bare-XOR enrichment with neutral signed counts, a more feed-forward profile; the antennal lobes (AL_L/R) are the only regions with significant signed *depletion*; and the gnathal ganglion (GNG), saddle (SAD), wedge (WED) and medulla (ME) show extreme enrichment of the signed motif. AL_R is a notable case: bare XOR is significantly enriched (induced *Z* = +2.63) while signed-XOR is significantly depleted (*Z* = −3.24). A caveat applies to all per-neuropil counts: the search is confined to each neuropil, so signed-XOR motifs whose feedback aggregators lie outside the neuropil (e.g. antennal-lobe projection neurons targeting the lateral horn or mushroom body) are not counted; the AL depletion may therefore partly reflect cross-neuropil feedback invisible to a within-neuropil search.

## C Mouse cortex: V1 biophysical model and MICrONS

### C.1 Allen Institute V1 biophysical model

#### Source data and extraction

The Allen Institute 2023 mouse V1 model [15, 34] distributes the connectome in the SONATA HDF5 format [35]: a `nodes.h5` file describing each neuron’s type and location, and `edges.h5` files listing synapses (source/target node identifiers, synapse-type identifier, numeric weight). The excitatory/inhibitory nature of a connection is implicit in the source neuron’s cell type (e.g. `i23Pvalb` sources emit inhibitory synapses). Our pipeline collapses this into a flat CSV with one row per synapse and columns `source_node_id`, `target_node_id`, `neuron_type`, `connection_type` (`EXC`/`INH`, from whether the source cell type begins with `e` or `i`), `syn_weight`. The result contains 7.0 10^7^ rows (5.5 10^7^ excitatory, 1.5 10^7^ inhibitory) across 230,922 neurons. Cell-type labels follow `[ei][layer][marker]` (markers: Pvalb, Sst, Htr3a, Rbp4, …); the inhibitory types used as role-6 hub candidates are listed in Table 8.

#### Weight filtering

The model assigns a continuous weight to every potential synapse; |*w*| spans more than twenty orders of magnitude, from ~ 10^−19^ (numerical residuals) to ~ 10^2^ (strong perisomatic Pvalb synapses). We retain only edges with |*w*| ≥ *w*_min_. All significance results below use the conservative *w*_min_ = 10^−1^ (≥100 *µ*V somatic perturbation, well above the cortical noise floor [36]), retaining strong, biologically-engaged synapses. The threshold is applied to |*w*|, not the raw signed weight: in the Allen model inhibitory weights lie in [−10, 0), so a raw-weight filter would silently discard every inhibitory edge.

#### Pivot-based significance

Whole-network randomisation of a ~ 4 × 10^7^-edge graph is computationally prohibitive, and the biologically relevant question is local: *is the signed-XOR over-represented around individual inhibitory neurons, and does this depend on cell type and layer?* For each inhibitory cell type and each randomly selected pivot *p* of that type, we (i) extract the *k*-hop (*k* = 2) bidirectional subgraph *G*_*p*_ around *p* at threshold *w*_min_, retaining all outgoing edges from *G*_*p*_ nodes so the counter can label boundary F^+^/F^−^ nodes correctly; (ii) count signed-XOR motifs with *p* in the role-6 position; (iii) generate *N* = 100 Roberts–Coolen null replicates of *G*_*p*_ (EXC and INH layers swapped independently, mixing = 100); and (iv) compute the same statistics as Section 3.4. Pivots were restricted to neurons with ≥ 2 outgoing inhibitory edges of strength |*w*| ≥ *w*_min_, with a fixed seed for reproducibility; the counter ran on 20 OpenMP threads. We report 60 pivots (~ 5 per inhibitory cell type) at *N* = 100 nulls each.

#### Results

Aggregating over 60 pivots (~5 per inhibitory cell type, *N* = 100 nulls each; Table 7) reveals a clear, biologically structured pattern. The signed-XOR is strongly over-represented in layers 2/3, 4 and 5 (per-pivot median folds 15–852×; single-pivot *Z* up to +7616; median empirical *p* at the floor 1*/*101 ≈ 0.0099), with a global enrichment of 315× (32,396,412 real motifs against 102,712 expected). Layer 4 dominates in absolute count, led by `i4Pvalb` (1.1 × 10^7^ motifs), with all three L4 types enriched 200–850-fold, two orders of magnitude above *Drosophila* AVLP_L. Layer 6 is *absent*: all three inhibitory types are non-significant (per-pivot |*Z*| < 2, *p* ≥ 0.13, real counts 0–357), with the null mean exceeding the real count in each case. The L6 result is a strong internal negative control: the same counter applied uniformly across all twelve types switches off cleanly at the L5/L6 boundary, where subgraph sizes are comparable (8,000–15,000 nodes, *>* 10^6^ edges), ruling out a size-driven combinatorial artefact. Within enriched layers the structure tracks known interneuron roles: by absolute count L4 is Pvalb-led (fast perisomatic feedforward gating [19, 20]) while its highest per-pivot fold is Htr3a; L2/3 is Htr3a-prominent (VIP-mediated disinhibitory selection [21, 22]); and L6 mediates corticothalamic feedback and global gain [37, 38], a regime where the local signed-XOR operation is not expected to dominate.

### C.2 MICrONS EM-reconstructed connectome

To test whether the layer structure of the Allen biophysical model is a property of the modelling assumptions or of cortical wiring itself, we applied the same motif-counting pipeline to the experimentally reconstructed MICrONS mouse V1 dataset [16]. Edge signs were assigned from cell-type predictions following Dale’s law (excitatory pyramidal sources positive, inhibitory interneuron sources negative), and MICrONS morphological interneuron classes were mapped onto the Allen Pvalb/Sst/Htr3a convention (basket cells to Pvalb, Martinotti cells to Sst, bipolar and neurogliaform cells to Htr3a). The cortical layer of each neuron is taken directly from the MICrONS cell-type label (the layer prefix of the excitatory subclass, e.g. `5P-IT` to layer 5, `6P-IT` to layer 6), rather than from a fixed soma-depth threshold.

Because the MICrONS segmentation is not exhaustively proofread, individual segments can carry merge artefacts that fuse two or more cells and inflate the apparent out-degree of the merged node. An out-degree ceiling is, however, not a safe way to remove them: a real layer-4 parvalbumin interneuron forms on the order of several thousand output synapses and therefore contacts of order several hundred distinct postsynaptic cells, so a low ceiling amputates genuine biological connectivity rather than artefacts. We confirmed this directly: capping the out-degree progressively removes the motif altogether (from 4.5 × 10^4^ motifs at a ceiling of 300 to zero below a ceiling of about 50), because it strips the dense but real connectivity that the motif requires, and the apparent pivot identity is unstable under the choice of ceiling. We therefore restrict the MICrONS analysis to the axon-proofread subset of The MICrONS Consortium [16], in which the out-degree of each source neuron is real by construction because its axon has been manually validated. This subset comprises 1,708 neurons and 86,793 signed edges after sign and layer assignment.

#### Results on the proofread subset

Table 9 reports the signed-XOR motif counts on the proofread MICrONS graph by inhibitory cell-type group and cortical layer. The qualitative layer structure of the biophysical model (Table 7) is reproduced in this measured, artefact-free connectivity. Layer 4 dominates the count (46.4% of all motifs, led by `i4Pvalb` with 55,870 motifs), layers 2/3 and 5 are intermediate (27.6% and 24.6%), and layer 6 contributes only 1.4% of motifs despite containing a fully populated complement of layer-6 interneurons (`i6Pvalb`, `i6Sst`, `i6Htr3a`) in the corrected assignment. The near-absence of the motif in layer 6 is therefore not a consequence of an empty layer, but a genuine paucity of the motif in a layer whose cells are present and available to form it. The parvalbumin pivot leads in every layer except layer 5, the single layer in which the Sst pivot exceeds it (`i5Sst` 23,369 versus `i5Pvalb` 15,467); this layer-5 Sst-prominence is consistent with the dynamical prediction of Section 5 (Prediction 1), in which the slow dendritic inhibition characteristic of layer-5 Sst interneurons realises the comparison by a route other than fast perisomatic veto.

#### On the null comparison

We report the MICrONS result as a description of the real wiring rather than as an enrichment over chance, because a degree- and cell-type-preserving null is not an adequate chance model for this substrate. On the proofread graph, the Roberts-Coolen edge-swap null that preserves each neuron’s per-sign degree and cell type produces *more* signed-XOR motifs than the real graph (of order 6 × 10^5^ versus 1.6 × 10^5^ across ten randomisations), not fewer. This is the expected behaviour of a degree-preserving null on a small, dense, spatially organised graph: rewiring that ignores distance and laminar position connects neuron pairs that the real wiring keeps apart, creating more motif opportunities than the structured biological graph contains. The real connectome is, if anything, more parsimonious in signed-XOR motifs than its degree-matched randomisation. We therefore do not claim enrichment of the motif over chance in MICrONS; establishing enrichment in a spatially embedded cortical graph requires a distance- and laminar-preserving null, which we identify as a priority follow-up. What the proofread MICrONS analysis does establish is that the motif is abundant and strongly layer-structured in measured cortical connectivity, with the same layer-4-dominant, parvalbumin-led, layer-6-sparse pattern seen in the Allen biophysical model.

#### Relation to the Allen model

The proofread MICrONS counts and the Allen-model counts (Table 7) agree on the principal qualitative features: a layer-4-dominant distribution, a parvalbumin-led pivot in layer 4, and a layer 6 that participates far less than the intracortical layers. They differ in the layer-5 share, which is substantially larger in the EM connectome (24.6%) than in the Allen model (1.3%); this may reflect either an underestimate of recurrent layer-5 connectivity in the biophysical model or genuine biology, and we flag it as a target for future investigation. We emphasise that the agreement concerns the layer-resolved structure of the motif, not an enrichment statistic: in the Allen model the motif is enriched against a degree-preserving null, whereas in the proofread MICrONS graph we make no enrichment claim for the reason given above.

## D Null model and randomiser

The null ensemble is generated by the unbiased degree-preserving edge-swap MCMC of Roberts and Coolen [17]. A swap selects two edges *a* → *b* and *c* → *d* within the same layer and rewires them to *a* → *d* and *c* → *b*, rejecting moves that would create a self-loop or a multi-edge. This preserves every node’s in-degree and out-degree within the layer exactly. The accept-all variant is used; Roberts and Coolen [17] show (their Sec. VI.C) that the resulting mobility bias is negligible for sparse fat-tailed biological networks. Default mixing is *T* = 100 proposed swaps per edge per layer; doubling to *T* = 200 does not measurably change motif counts.

Swaps occur *within* sign layers, so a neuron emitting only inhibitory edges in the real network emits only inhibitory edges in every null. For *Drosophila* the layers are the predicted neurotransmitters (ACH, GABA, GLUT randomised independently). For *C. elegans* the layers are the curated excitatory and inhibitory classes (all excitatory edges, including the aminergic ones curated EXC, swap within one pool, all inhibitory within another), which is the correct Dale-preserving null for a connectome whose functional sign is curated per edge rather than implied by neurotransmitter. For mouse V1 the layers are the cell-type-derived EXC and INH classes of the local subgraph; for the MICrONS analysis the layering is finer still, with one layer per mapped cell type (Section C).

### Availability

The counters, the randomiser, the bash significance drivers, synthetic planted-motif test cases, and the run logs are at https://github.com/jesusmarcodelucas/signed-xor-motifs [12].

